# Transcriptional control of motor pool formation and motor circuit connectivity by the LIM-HD protein Isl2

**DOI:** 10.1101/2022.11.07.515417

**Authors:** Yunjeong Lee, In Seo Yeo, Namhee Kim, Dong-Keun Lee, Kyung-Tai Kim, Jiyoung Yoon, Young Bin Hong, Byung-Ok Choi, Yoichi Kosodo, Daesoo Kim, Jihwan Park, Mi-Ryoung Song

**Affiliations:** School of Life Sciences, Gwangju Institute of Science and Technology, Oryong-dong, Buk-gu, Gwangju, 61005, Republic of Korea; Fermentation Regulation Technology Research Group, World Institute of Kimchi, Gwangju 61755, Republic of Korea; Jeonbuk Department of Inhalation Research, Korea Institute of Toxicology, Jeongeup-si, Jeollabuk-do, Republic of Korea; Korea Brain Research Institute, 61, Cheomdan-ro, Dong-gu, Daegu, 41062, Republic of Korea; Department of Neurology, Samsung Medical Center, Sungkyunkwan University School of Medicine, Seoul, 06351, Republic of Korea; Department of Biochemistry, College of Medicine, Dong-A University, Busan, Republic of Korea; Department of Biological Sciences, Korea Advanced Institute of Science and Technology (KAIST), Daejeon, Republic of Korea

## Abstract

The fidelity of motor control requires the precise positional arrangement of motor pools and the establishment of synaptic connections between these pools. In the developing spinal cord, motor nerves project to specific target muscles and receive proprioceptive input from the muscles via the sensorimotor circuit. LIM-homeodomain transcription factors are known to successively restrict specific motor neuronal fates during neural development; however, it remains unclear to what extent they contribute to limb-based motor pools and locomotor circuits. Here, we showed in mice that deletion of *Isl2* resulted in scattered motor pools, primarily in the median motor column and lateral LMC (LMCl) populations, and lacked Pea3 expression in the hindlimb motor pools, accompanied by reduced terminal axon branching and disorganized neuromuscular junctions. Transcriptomic analysis of *Isl2*- deficient spinal cords revealed that a variety of genes involved in motor neuron differentiation, axon development, and synapse organization were downregulated in hindlimb motor pools. Moreover, the loss of *Isl2* impaired sensorimotor connectivity and hindlimb locomotion. Together, our studies indicate that *Isl2* plays a critical role in organizing motor pool position and sensorimotor circuits in hindlimb motor pools.

## Introduction

The establishment of motor circuits relies on the stereotyped nuclear organization of motor neurons, termed motor columns, and motor pools at distinct co-ordinates that innervate discrete muscles (Romanes, 1964; Vanderhorst & Holstege, 1997). Anatomically defined motor columns reside in the spinal cord along the rostrocaudal axis. Within each columnar group, motor neurons are further subdivided into motor pools. Many transcription factors and signals that contribute to the biological processes in which motor neurons diverge and differentiate into distinct functional subtypes have been identified: spatiotemporal generation of motor columns and motor pools, long-range axon projection to the target muscles, and establishment of elaborate axonal and dendritic arbors for synaptic connections. For instance, the combinatorial expression of LIM-homeodomain (HD) transcription factors specifies pan-motor neuron identity, which further diverges into motor columns across multiple segments, including the median motor column (MMC), the hypaxial motor column (HMC), the lateral motor column (LMC), and the preganglionic column (PGC) neuronal fates (Pfaff, Mendelsohn, Stewart, Edlund, & Jessell, 1996; Tsuchida et al., 1994). These transcription factors subsequently orchestrate a variety of motor neuron differentiation programs, such as cell migration, axonal pathfinding, and target specificity.

Simultaneously, the arrangement of motor columns and motor pools along the rostrocaudal axis is shaped by Foxp1, the expression for which is dictated by Hox transcription factors that employ positional cues along the body axis; Foxp1, alongside Hox transcription factors, induces LMCs at brachial and lumbar levels, and PGCs at the thoracic level(Dasen, De Camilli, Wang, Tucker, & Jessell, 2008; Dasen, Tice, Brenner-Morton, & Jessell, 2005; Rousso, Gaber, Wellik, Morrisey, & Novitch, 2008). Individual motor pools are further subdivided and matured with the help of E26 transformation-specific (ETS) transcription factors, including *Pea3* and ETS-related protein 81 (ER81), which contribute to the assembly of sensorimotor circuits (Arber, Ladle, Lin, Frank, & Jessell, 2000; Livet et al., 2002; Vrieseling & Arber, 2006). Thus, a variety of players in series contribute to the entire collection of motor pools in place and ensure their connectivity and functions (Dasen et al., 2008; Rousso et al., 2008; Surmeli, Akay, Ippolito, Tucker, & Jessell, 2011).

Given that more than 50 muscles in the limb of most tetrapods are innervated by distinct limb motor pools, there likely exist a variety of molecular features that shape individual motor pools during spinal cord development (Sullivan, 1962). In particular, several lines of evidence have suggested that some designated molecular players construct hindlimb motor pools. The discovery of neuromesodermal progenitors specialized for building the caudal part of the spinal cord indicates that the origin of caudal motor neurons is different from that of rostral motor neurons (Henrique, Abranches, Verrier, & Storey, 2015; Shaker et al., 2021). For instance, the fate mapping study of NMP populations revealed a rostral-to-caudal ascending gradient of motor neuron generation in the developing spinal cord, which indicated that a distinct genetic program guides lumbar motor pools separately (Shaker et al., 2021). Additionally, some *Hox* genes, alongside their initial role in body patterning, are selectively expressed in motor neuron-lineage cells during the postmitotic stages and further define their caudal segmental identity. Interventions of caudal *Hox* genes such as *Hoxd9*, *Hoxc10*, and *Hoxd10* often lead to topographic disorganization and inappropriate targeting of LMC MNs at the lumbar level (de la Cruz, Der-Avakian, Spyropoulos, Tieu, & Carpenter, 1999; Y. Wu, Wang, Scott, & Capecchi, 2008). However, fine-tuning of the three-dimensional organization of individual motor pools may not be sufficiently addressed solely by *Hox* networks because the elongated shape of motor pools sometimes encompasses several spinal cord segments, and *Hox* genes mainly assign positional information along the rostrocaudal axis rather than the dorsoventral and mediolateral axes. Furthermore, additional players known to act together with or be under the control of *Hox* genes to impose LMC fates appear to be valid for both limb segments. Foxp1 activity induces LMC identity regardless of their limb level; both brachial and lumbar motor neurons are equally affected in the absence of Foxp1 and possess scattered and misspecified LMC neurons (Dasen et al., 2008). Retinoid receptor signaling or one-cut factors also induce LMC identity at both limb segments (Francius & Clotman, 2014; Sockanathan, Perlmann, & Jessell, 2003). Similarly, ETS factors exhibit limited expression on LMC motor pool subsets at both limb levels, and the loss of these factors, such as *Pea3* or *Er81*, leads to the defective organization of several LMC motor pools with reduced dendritic arborization and impaired neuromuscular junction formation (mostly examined at the brachial level) (Livet et al., 2002; Vrieseling & Arber, 2006). Thus, their potential role in lumbar motor pools remains unexplored. Finally, recent genome-wide transcriptomic studies at the single-cell level have identified extensive molecular signatures for individual motor columns, further expanding the segmental diversity of spinal motor neurons (Amin et al., 2021; Delile et al., 2019; Liau et al., 2021; W. Wang, Cho, Lee, & Lee, 2022). However, detailed comparisons among LMC populations of the two limb segments and the identification of a genetic program that elicits the arrangement of hindlimb motor pools remain elusive.

Isl1 and Isl2 are two related LIM-HD transcription factors that shared approximately 75% of their identity and are highly expressed in postmitotic motor neurons in an overlapping manner. Genetic studies in mice and zebrafish have revealed that the contribution of *Isl1* is significant, perhapse more cruicial than that of *Isl2*, demonstrating that the removal of *Isl1* is sufficient to induce defects in motor neuronal fates, axonal navigation, neurotransmitter identity, and electrical excitability (Liang et al., 2011; Moreno & Ribera, 2014; Pfaff et al., 1996; Wolfram, Southall, Brand, & Baines, 2012). However, several lines of evidence have supported the idea that *Isl2* is equally important for motor neuron development. First, in *Isl2*-null mice, thoracic motor neurons are affected, reflected by scattered or misspecified motor neuron subsets (Thaler et al., 2004). Similarly, misexpression of the dominant negative form of *Isl2* or knockdown (KD) of *Isl2* in zebrafish results in mispositioned motor neuron cell bodies and defective axon growth (Hutchinson & Eisen, 2006; Segawa et al., 2001). Further, successful rescue experiment on *isl1* morphants using *isl2* mRNA indicates that both islet proteins can be equally potent in neurons (Hutchinson & Eisen, 2006). Second, although detailed expression requires further investigation, *Isl2*, similarly to *Isl1*, is also broadly expressed in motor neurons during motor neuron development (Thaler et al., 2004). Specifically, LMCl is a population that expresses *Isl2* but not *Isl1*, which raises the possibility that *Isl2* plays an independent role in LMC1, besides what has been reported in thoracic motor neuron subsets. Lastly, recent advances in transcriptomic analysis have allowed us to visualize more detailed spatiotemporal changes in the gene expression of islet factors during motor neuron development, where expression persists until the postnatal period (Amin et al., 2021; Catela, Chen, Weng, Wen, & Kratsios, 2022; Delile et al., 2019; Rayon, Maizels, Barrington, & Briscoe, 2021). However, the substantial overlap between the expression of *Isl1* and *Isl2* during various stages of differentiating motor neurons has only allowed limited interpretations. Overall, despite considerable similarities or redundancies in islet factors, the role of *Isl2* in the context of motor neuronal development has been relatively underestimated and warrants further investigation.

In this study, we demonstrated that Isl2 is enriched in proximal motor pools expressing Pea3 and is involved in setting the position of their cell bodies directing the assembly of the motor neuronal circuit, particularly at the hindlimb level. *Isl2*-null mice displayed defective hindlimb movements, which correlated strongly with disorganized neuromuscular junction formation and reduced axonal and dendritic arborization of motor neurons. Genome-wide transcriptomic analysis revealed that *Isl2* acts as a master regulator of various genes involved in the development and differentiation of particular motor pools located in the hindlimb. Our results suggest that LIM-HD transcription factors, together with Hox and ETS transcription factors, fine-tune the organization of hindlimb motor pools.

## Results

### Spatiotemporal expression of islet transcription factors during motor neuron development

Islet transcription factors *Isl1* and *Isl2* are expressed immediately after motor neuron generation; however, the detailed expression of islet transcription factors during motor neuron development at the cellular level remains uncertain. Thus, we used published single-cell RNA-sequencing data obtained from E12 motor neurons; sorted from the cervical to lumbar regions of the spinal cord; and reanalyzed gene expression profiles, focusing on the changes in expression levels of islet transcription factors (Amin et al., 2021). Approximately 3,483 cells of developing motor neurons were divided into eight subgroups representing major motor columns, such as pMN, immature, MMC, P/HMC, LMCm, LMCl, PGCa, and PGCb (Supplementary Figure 1a and c). Uniform Manifold Approximation and Projection (UMAP) revealed the progression of pMN progenitors into immature motor neurons, MMC, P/HMC, LMCm, LMCl, PGCa, and PGCb. We observed high expression of *Isl1* in earlier immature motor neurons and slightly lower expression in most differentiated motor neurons, including MMC, P/HMC, LMCm, and PGCa, but not LMCl, which did not express *Isl1*. However, *Isl2* expression arose later than *Isl1* in immature motor neurons, and its expression continued to persist in MMC and P/HMC neurons. Low but even expression of *Isl2* was observed in most LMC neurons, while a few PGCa and b cells expressed *Isl2* (Supplementary Figure 1b). Differential expression of *Isl1* and *Isl2* between different motor columns was also observed in E12.5 mouse spinal cords (Supplementary Figure 1d). Thus, both islet transcription factors may serve overlapping and nonredundant functions in differentiating motor neurons, and the contribution of *Isl2* may be more significant among MMC and LMCl populations, in which *Isl2* expression is more pronounced than *Isl1* expression.

### Settling position of motor neurons is impaired in the absence of *Isl2*

Because *Isl2* is broadly expressed in postmitotic motor neurons, we next examined the overall organization of motor columns in *Isl2*-null mice. Given the panmotor neuronal expression of *Isl2* across the multiple segments of the spinal cord, it is likely that additional motor neuronal populations would be similarly affected in the absence of *Isl2*. The expression of various motor neuronal markers was examined in adjacent sections: *Raldh2* and Foxp1 (LMC markers), Hb9 (MN marker), and Lhx3 (MMC marker). At E13.5, the number of MMC and LMC neurons at the limb level remained unchanged (Figure 1a–e and Supplementary Table S2). At the thoracic level, there were some changes in the number of MMC and PGC neurons and defective PGC neurons, with a lower expression of nitric oxide synthase (NOS) (Figure 1a, f, and g) (Thaler et al., 2004). Notably, we found that motor neuronal cell bodies were slightly scattered and occupied the region between the MMC and LMC at the limb level, which was maintained until adulthood (Figure 1a, h, and i). The ectopic cells at the lumbar level consisted of Lhx1^+^ LMCl (52%), Lhx3^+^ MMC neurons (29%), and unidentified Hb9::GFP-expressing cells. Mislocalized motor neurons remained in the adult *Isl2*-null lumbar spinal cord with a similar number of motor neurons, which collectively indicated that the position of the MMC and LMCl was affected in the absence of *Isl2* (Figure 1h and i and Supplementary Figure 2).

**Figure 1.**
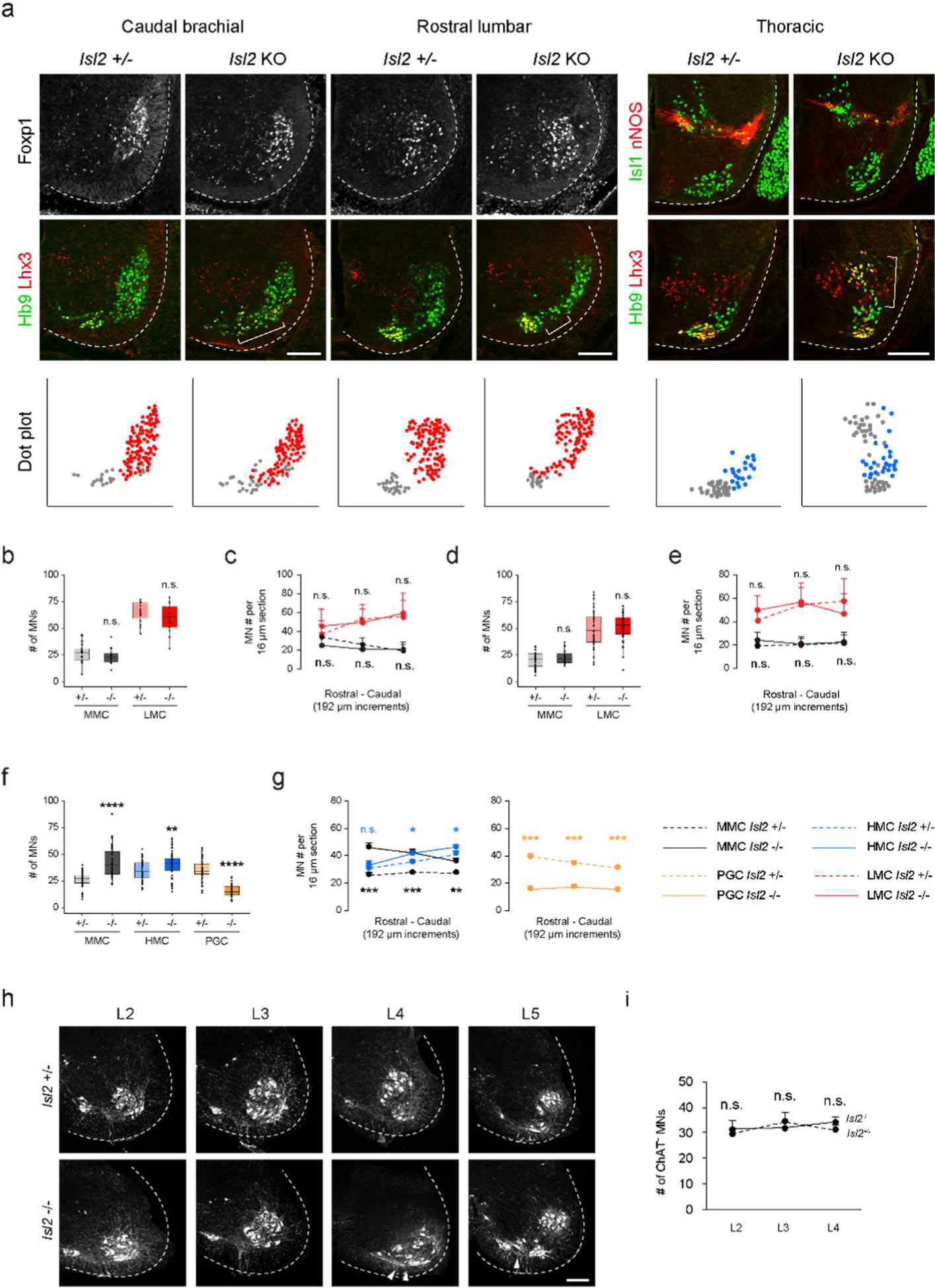
*Isl2*-null mice had scattered spinal motor neurons with disorganized motor columns. (a) Expression of Foxp1, Hb9, Lhx3, Isl1 and nNOS in E13.5 mouse embryonic spinal cords as indicated. Brackets mark ectopic motor neurons in *Isl2-*null mice. For spatial plots, MMC (gray, Lhx3^+^Hb9^+^) neurons, LMC (red, Foxp1^+^) neurons, and PGC (blue, Isl1^+^ Hb9^+^Lhx3^−^) neurons from the representative images above are plotted. (b, d, f) Average number of motor neurons in 16 μm transverse sections of the MMC, LMC, and HMC in caudal brachial (b), rostral lumbar (d), and thoracic (f) spinal segments. Box plots showing feature distribution represent median (center line), first and third quartiles (box boundaries), and 10th and 90th percentiles (whisker). (c, e, g) The number of motor neurons was counted in transverse sections at 192 μm intervals in caudal brachial (c), rostral lumbar (e), and thoracic (g) spinal cords. (h, i) Representative images and quantification of adult adjacent lumbar spinal cords of *Isl2* +/− and *Isl2* KO *Hb9::GFP* mice. Ectopic motor neurons are labeled with arrowheads (three animals per group; SEM is shown for c, e, g, i ; unpaired Student’s t-test; see Supplementary Table S2 for detailed n and statistics). \**p* < 0.05, \*\**p* < 0.01, \*\*\**p* < 0.001, \*\*\*\**p* < 0.0001, n.s = not significant. Scale bars, 100 μm.

To further characterize the scattered motor neurons at the limb level, the organization of motor pools at the brachial level was examined according to the combinatorial pattern of markers for the motor columns and pools, such as Hb9::GFP, Foxp1, Isl1, Pea3, and Scip, in consecutive transverse spinal cord sections (Figure 2). A previous study reported that the organization of the Hb9^+^Lhx3^+^ MMC population is disrupted at the thoracic level when *Isl2* is absent (Thaler et al., 2004). We similarly found that MMC neurons were scattered dorsolaterally and located ectopically, either within the LMC columns or immediately dorsal to the motor columns in *Isl2-*deficient brachial spinal cords (Figure 1a). However, the segregation between the LMCm and LMCl was relatively spared when determined by Isl1 and Foxp1 immunoreactivity (Figure 2a and b). Furthermore, the distribution of major brachial motor pools, defined by Pea3 and Scip expression, remained unchanged in *Isl2* mutant mice. At the C6 to C8 level, the clustering of Pea3-expressing motor pools innervating Isl1^+^, Foxp1^+^, Pea3^+^, and GFP^low^ cutaneous maximus (CM) and Isl1*^−^*, Foxp1^+^, Pea3^+^, and GFP^high^ latissimus dorsi (LD) muscles, and Scip-expressing motor pools innervating Foxp1^+^, Scip^+^, and GFP^+^ flexor carpi ulnaris (FCU), was unchanged in *Isl2* mutant mice (Figure 2a, b, d) (Dasen et al., 2005; Livet et al., 2002). In line with this result, the number of individual motor pools was also unchanged in *Isl2* mutant mice (Figure 2c). Therefore, at the brachial level, MMC neurons were mostly scattered, whereas the position of other populations remained relatively normal.

**Figure 2.**
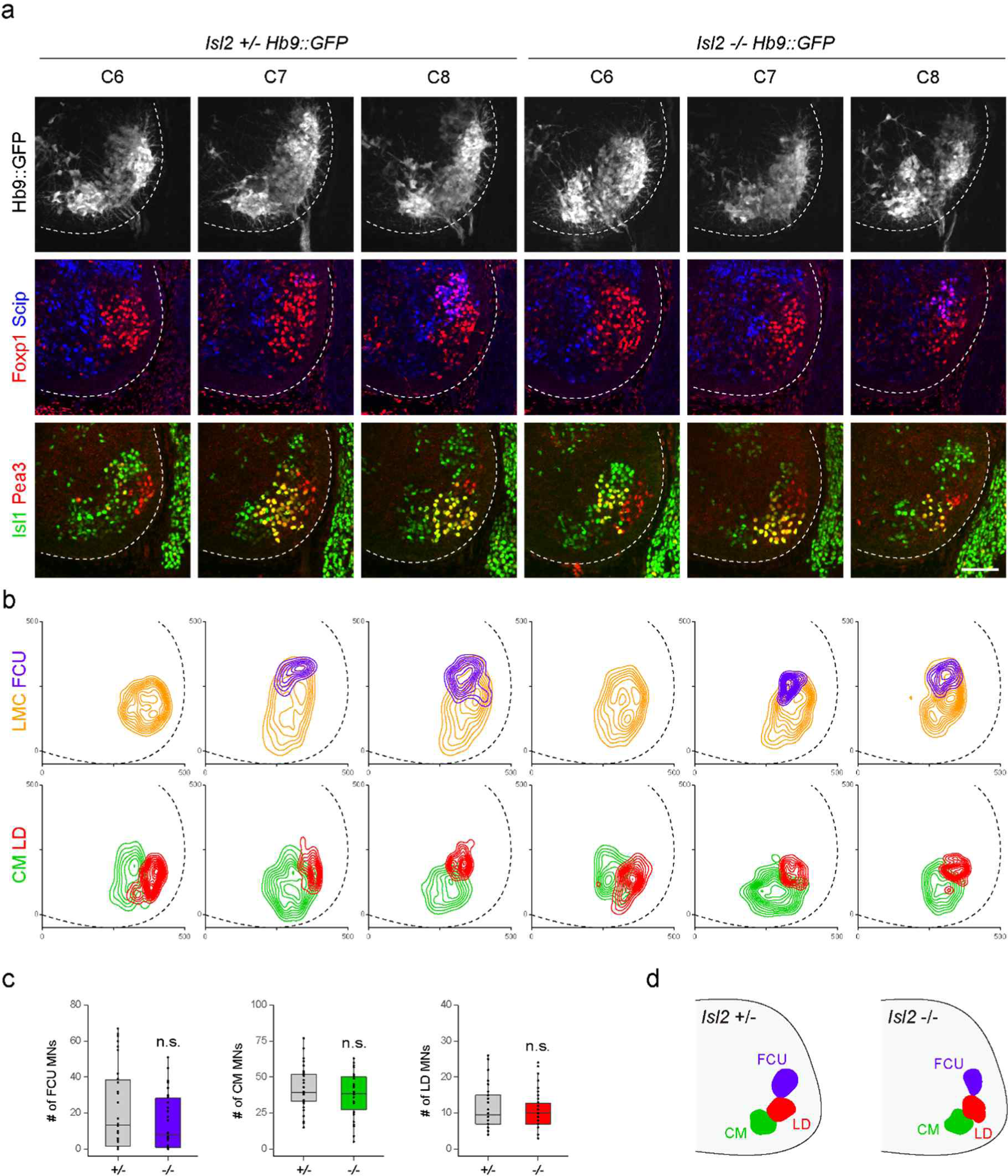
Distribution of LMC motor pools in E13.5 control and *Isl2* mutant brachial spinal cords. (a) Representative images and spatial plots of motor neurons labeled with Hb9-GFP, Foxp1, Scip, Isl1, and Pea3 in adjacent sections of C6 to C8 spinal cords. Triple-labeled images show Foxp1, Scip, and Hb9::GFP expression in the same section. (b) Contour plots for Foxp1^+^Hb9::GFP^+^ LMC neurons (orange); Foxp1^+^Scip^+^, GFP^+^ FCU (purple); Isl1^+^Foxp1^+^Pea3^+^, GFP^low^ CM (green); and Isl1^−^Foxp1^+^Pea3^+^, GFP^high^ LD (red). (c) The number of cells in FCU, CM, and LD motor pools (three animals per group; unpaired Student’s t-test; see Supplementary Table S2 for detailed n and statistics). Box plots showing feature distribution represent median (center line), first and third quartiles (box boundaries), and 10th and 90th percentiles (whisker). n.s = not significant. (d) Summary diagram shows the position of major brachial motor pools. Scale bar, 100 μm.

Next, we reconstructed the detailed motor pool distribution at the lumbar level by analyzing the expression of motor neuronal markers, Hb9::GFP, Foxp1, Lhx3, Nkx6.1, Lhx1, Pea3, Scip, and *Er81* in consecutive transverse spinal cord sections from the rostral lumbar level (Figure 3). The lumbar LMC consists of multiple motor pool clusters, which can be identified by their stereotyped position and distinct marker expression: Isl1^+^, Foxp1^+^, Hb9^+^, and GFP^low^ hamstring (H); Nkx6.1^−^, *Er81*^+^, Lhx1^+^, and GFP^high^ vastis (V); Lhx1^+^, Foxp1^+^, Pea3^+^, and GFP^high^ rectus femoris/tensor fasciae latae (Rf/Tfl); Isl1^+^, Foxp1^+^, Hb9^+^, and GFP^low^ gastrocnemius (Gs); Nkx6.1^+^, Lhx1^+^, Foxp1^low^, *Er81*^−^, Pea3^−^, and GFP^high^ tibialis anterior (Ta); and Nkx6.1^-^, Lhx1^+^, Foxp1^+^, Pea3^+^, and GFP^high^ gluteus (Gl) (Arber et al., 2000; De Marco Garcia & Jessell, 2008; Greene, 1935; Landmesser, 1978; Livet et al., 2002; McHanwell & Biscoe, 1981; Surmeli et al., 2011). Overall, the positions of the Hb9^+^ GFP^+^ Lhx3^+^ MMC and Hb9^+^ GFP^+^ Foxp1^+^ LMC were relatively intact; however, the distributions of Nkx6.1 and Lhx1 within the LMC were less distinct in *Isl2* mutant mice (Figure 3a, b, d, Supplementary Figure 3). The clustering of H, Gs, and Ta pools was relatively normal in *Isl2* mutant mice, whereas some *Er81*^+^ were slightly scattered laterally, and Nkx6.1^+^ Ta motor neurons were scattered anteriorly in the L2 segment. The number of *Er81*^+^ and Ta motor neurons was unchanged (Figure 3c). Remarkably, the number of Pea3-expressing cells including the Rf/Tfl and Gl pools in the L4 segment was significantly downregulated whose individual position and overall distribution were shown in dot and contour plots (Figure 3a-c and Supplementary Figure 3). Taken together, MMC neurons at all levels and some LMC motor pools at the lumbar level were dispersed in the absence of *Isl2*.

**Figure 3.**
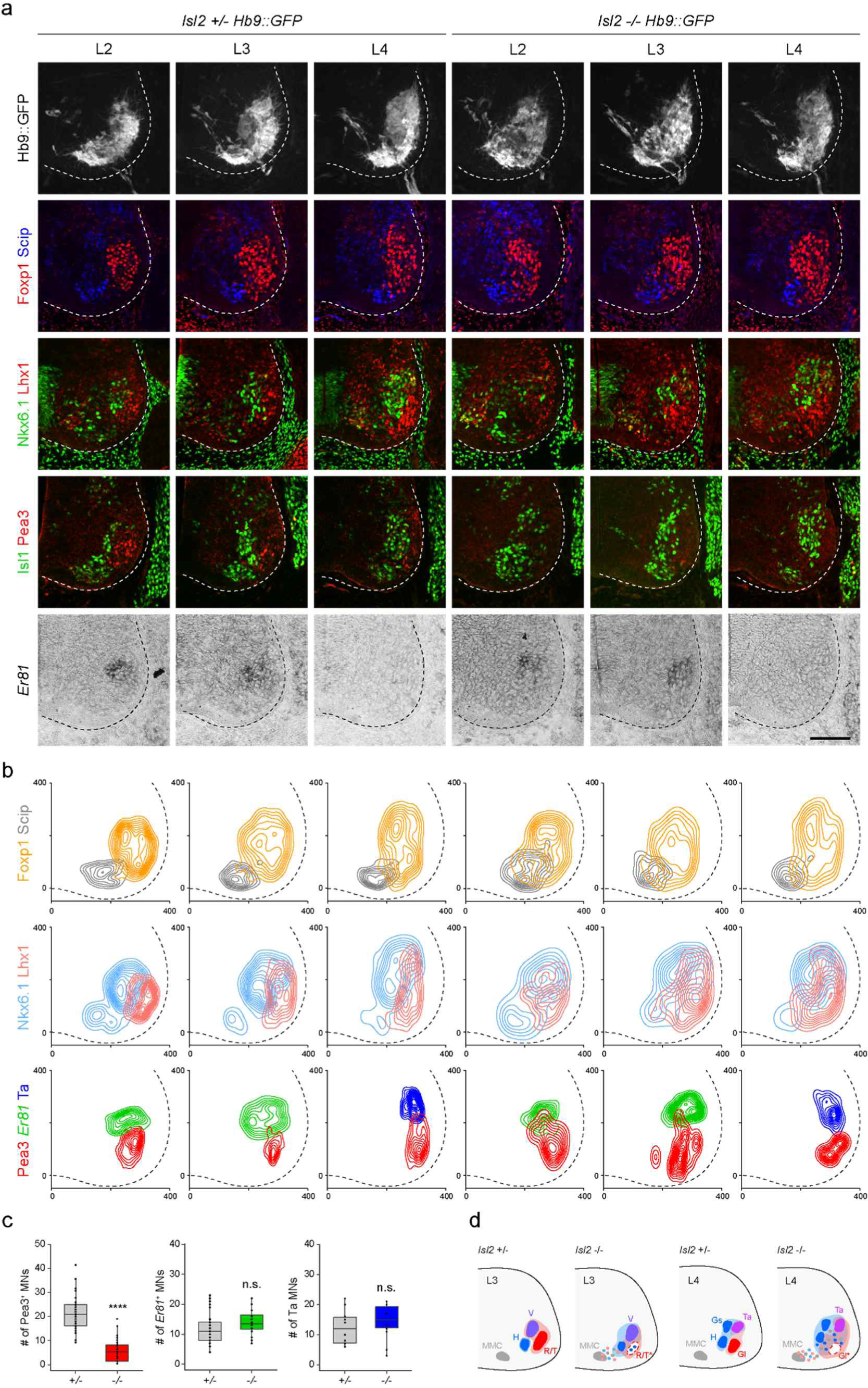
Motor pool position in E13.5 control and *Isl2* mutant lumbar spinal cords. (a, b) Representative images and spatial plots of motor neurons labeled with Foxp1, Lhx3, Nkx6.1, Lhx1, Scip, Pea3, and *Er81* in adjacent sections of L2 to L4 spinal cords. Triple-labeled images show Foxp1, Lhx3, and Hb9::GFP expression in the same section. Contour plots for Lhx3^+^, Hb9::GFP^+^ MMC neurons (gray); Foxp1^+^, Hb9::GFP^+^ LMC neurons (orange); Nkx6.1^+^, Foxp1^+^ LMCm neurons (light blue); Lhx1^+^, Hb9::GFP^+^ LMCl neurons (pink); Pea3 (red); *Er81* (green); and Nkx6.1^+^, Lhx1^+^ Ta (blue) neurons were generated from adjacent sections of the same embryo (n = 4 adjacent sections, see Supplementary Table S2 for detailed n). (c) The number of neurons in LMCm, LMCl, Pea3, *Er81*, and Ta motor pools (three animals per group; unpaired Student’s t-test; see Supplementary Table S2 for detailed n and statistics). Box plots showing feature distribution represent median (center line), first and third quartiles (box boundaries), and 10th and 90th percentiles (whisker). n.s = not significant. \*\*\*\**p* < 0.0001. (d) Summary diagram shows distributions of the hamstring (H), vastus (V), rectus femoris (R), tensor fascia latae (T), gastrocnemius (Gs), tibialis anterior (Ta), and gluteus (Gl) motor pools. Motor pool position from (Surmeli et al., 2011; Vanderhorst & Holstege, 1997). Asterisks indicate misspecified motor pools. Scale bar, 100 μm.

### Isl2 induces *Pea3* transcripts in lumbar motor pools

Scattered motor neurons and the absence of Pea3 expression in *Isl2* mutant lumbar motor pools led us to reason that Isl2 activates *Pea3* expression because its absence also resulted in positional defects in MNs (Livet et al., 2002). Thus, we examined Pea3 expression in *Isl2* mutant mice during the segregation of limb motor pools from E11.5 to E13.5. We used two mouse lines, *Isl2* KO mice and *Isl2* conditional knockout (cKO) mice, which were generated by crossing *Isl2^flox/flox^* mice with *Isl2^+/−^*; Olig2-Cre mice (Supplementary Figure 4a and b) (Dessaud et al., 2007; Kong et al., 2015). *In situ* hybridization analysis showed that the transcript level of *Isl2* was specifically downregulated among motor neurons not in others (Supplementary Figure 4c). Immunohistochemistry and RT-PCR analysis using spinal cord tissues revealed that both the proteins and transcripts of *Pea3* were selectively downregulated in lumbar motor pools but not in brachial motor pools of *Isl2* KO and *Isl2* cKO spinal cords (Figure 4a and b and Supplementary Figure 5).

**Figure 4.**
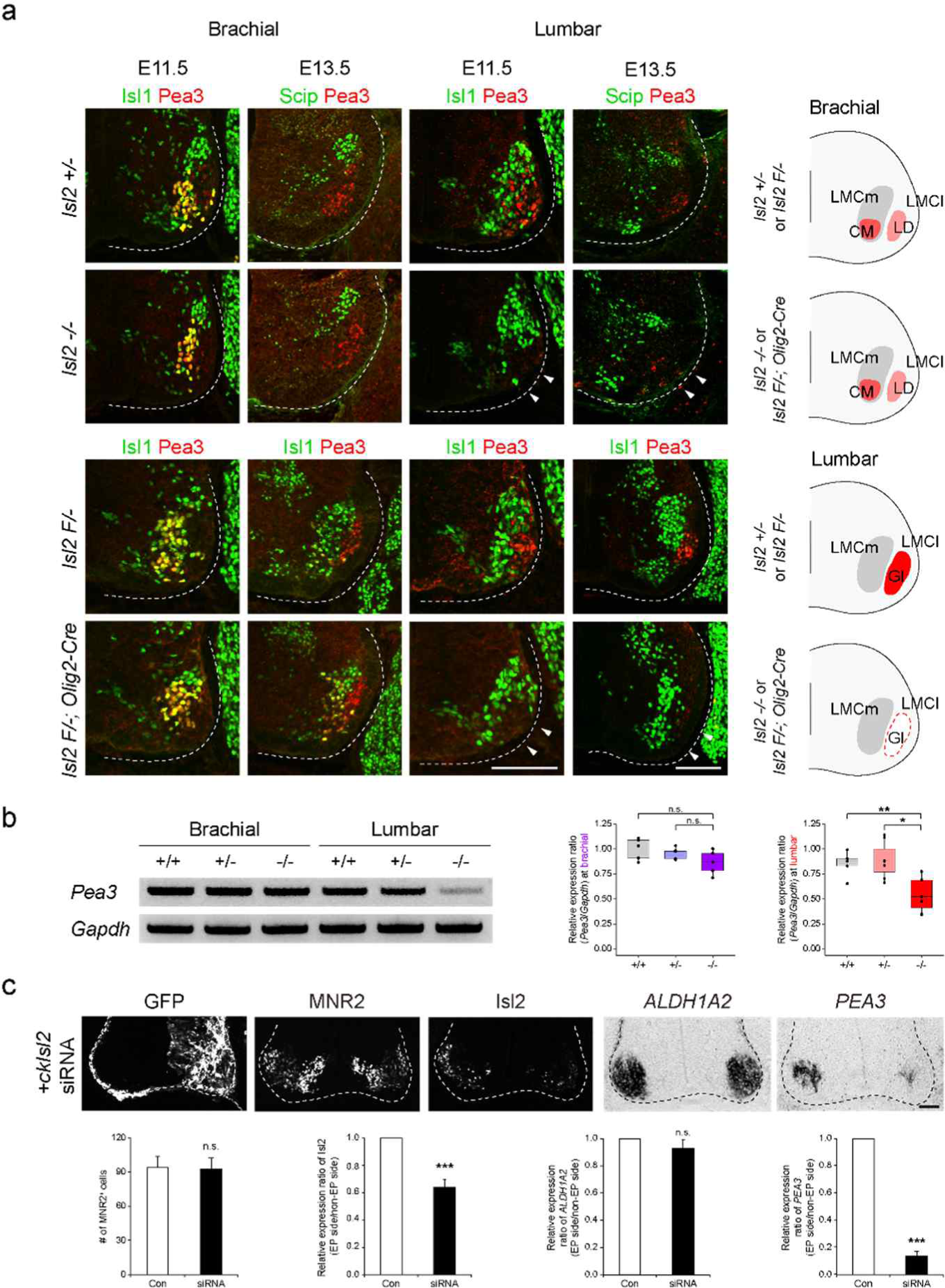
The elimination of Pea3 expression in *Isl2*-null motor neurons. (a) Expression of Isl1, Scip, and Pea3 in E11.5 and E13.5 brachial and lumbar spinal cords of *Isl2 +/*−, *Isl2 KO*, *Isl2 F/*−, and *Isl2 F/*−; Olig2-Cre animals. Note that Pea3 expression disappeared in lumbar but not brachial motor neurons (arrowheads). Summary diagram shows the position of LMC motor pools innervating the cutaneous maximus (CM), latissimus dorsi (LD), and gluteus (Gl) muscles from each genotype. Scale bars, 100 μm. (b) RT-PCR results and quantification show lower levels of *Pea3* transcript expression in E12.5 *Isl2-*null lumbar spinal cords (n = 3 animals per genotype; box plots showing feature distribution represent median (center line), first and third quartiles (box boundaries), and 10th and 90th percentiles (whisker); one-way ANOVA with Tukey’s test; see Supplementary Table S2 for detailed n and statistics). \**p* < 0.05, \*\**p* < 0.01, n.s = not significant. (c) Expression of GFP, MNR2, Isl2, *ALDH1A2*, and *PEA3* in HH stage 25 chick neural tubes, electroporated with siRNA against *Isl2*. Quantification of MNR2-expressing motor neurons, fluorescent intensity of Isl2 expression, and relative expression of *ALDH1A2* and *PEA3*. Note that the *PEA3* transcript was downregulated on the electroporated side (right; n = 5 animals per group; SEM is shown; see Supplementary Table S2 for detailed n and statistics). \*\*\**p* < 0.001, n.s = not significant. Scale bars, 50 μm.

To test whether acute downregulation of *Isl2* is sufficient to abolish *Pea3* transcription, we knocked down *Isl2* expression in the chick neural tube by siRNA. Efficient KD of *Isl2* was tested in a cell line misexpressing HA-tagged ckIsl2 (Supplementary Figure 6). When *Isl2* siRNA was introduced into the chick neural tube by in ovo electroporation, the number of Isl2-expressing cells and the intensity of Isl2 in these cells were reduced, while the total number of MNR2-labeled motor neurons remained unchanged (Figure 4c). Consistent with the results in mice, the *PEA3* transcript level was significantly downregulated in *Isl2-*KD motor neurons (Figure 4c). Collectively, these results suggest that Isl2 is necessary for gene transcription of *Pea3* in motor neurons in a cell-autonomous manner, which may correlate with the correct positioning of motor pools.

### Transcriptomic analysis to define genes is associated with the development of lumbar motor pools

Because the motor pools at the lumbar level were more severely affected in the absence of *Isl2* compared with those at other axial levels, we sought to identify sets of genes under the control of *Isl2*, particularly at the lumbar segment. We applied two criteria to search for the genes responsible. First, we focused on downregulated genes because Isl2 is expected to act as an activator similar to Isl1 (S. Lee et al., 2008; S. K. Lee & Pfaff, 2003). Second, by comparing DEGs at the brachial and lumbar levels, we selected genes that were only downregulated at the lumbar level. We conducted bulk RNA-seq analysis using E12.5 brachial and lumbar ventral spinal cords. To obtain motor neuron-enriched tissues, we dissected the ventral spinal cord from an open-book preparation. The segmental identity of the spinal cord was verified according to the expression of the correct *Hox* code at each axial level; *Hoxc5*, *Hoxc6*, and *Hoxc8* transcript levels were upregulated in the brachial spinal cord, and *Hoxd9*, *Hoxa10*, *Hoxc10*, *Hoxd10*, *Hoxa11*, *Hoxc11*, and *Hoxd11* transcript levels were upregulated in the lumbar spinal cord (Supplementary Figure 7g) (Hayashi et al., 2018). The differentially expressed gene (DEG) analysis between *Isl2* KO and *Isl2* heterozygote mice revealed that 140 genes and 159 genes, respectively, were downregulated at the brachial and lumbar levels (<−0.5 log_2_-FC, raw *p* < 0.5). We then collected DEGs that overlapped with the list of genes from the scRNA-seq dataset obtained from embryonic motor neurons to further identify motor neuron-specific genes (Amin et al., 2021). As a result, 44 and 87 DEGs were specifically downregulated at the brachial and lumbar levels of *Isl2* KO spinal cords, respectively, whereas nine DEGs were commonly downregulated in both segments (Figure 5a–e and Supplementary Table S3). Notably, gene ontology (GO) analysis revealed that the downregulated genes at the lumbar level in *Isl2*-null embryos were predominantly subcategorized into the neuropeptide signaling pathway, synaptic signaling, behavior, regulation of ion transport, neuron differentiation, and axon development (Figure 5g). In contrast, at the brachial level, no categories involving synapse organization, axon development, and neuron differentiation were detected (Figure 5f). When the expression of DEGs was validated in *Isl2* mutant spinal cords, the expression of *A730046J19Rik*, *Axna2*, *Chodl*, Etv4 (Pea3), *Prph*, *Kcnab1*, *C1ql3*, and *Hoxc11* was markedly downregulated in the most ventrolaterally located motor pool subsets at the E12.5 lumbar spinal cords, which is consistent with normalized read count values (Figure 5h). In addition, nNOS/Nos1 immunoreactivity was downregulated in PGC neurons, as reported previously (Figure 5h) (Thaler et al., 2004).

**Figure 5.**
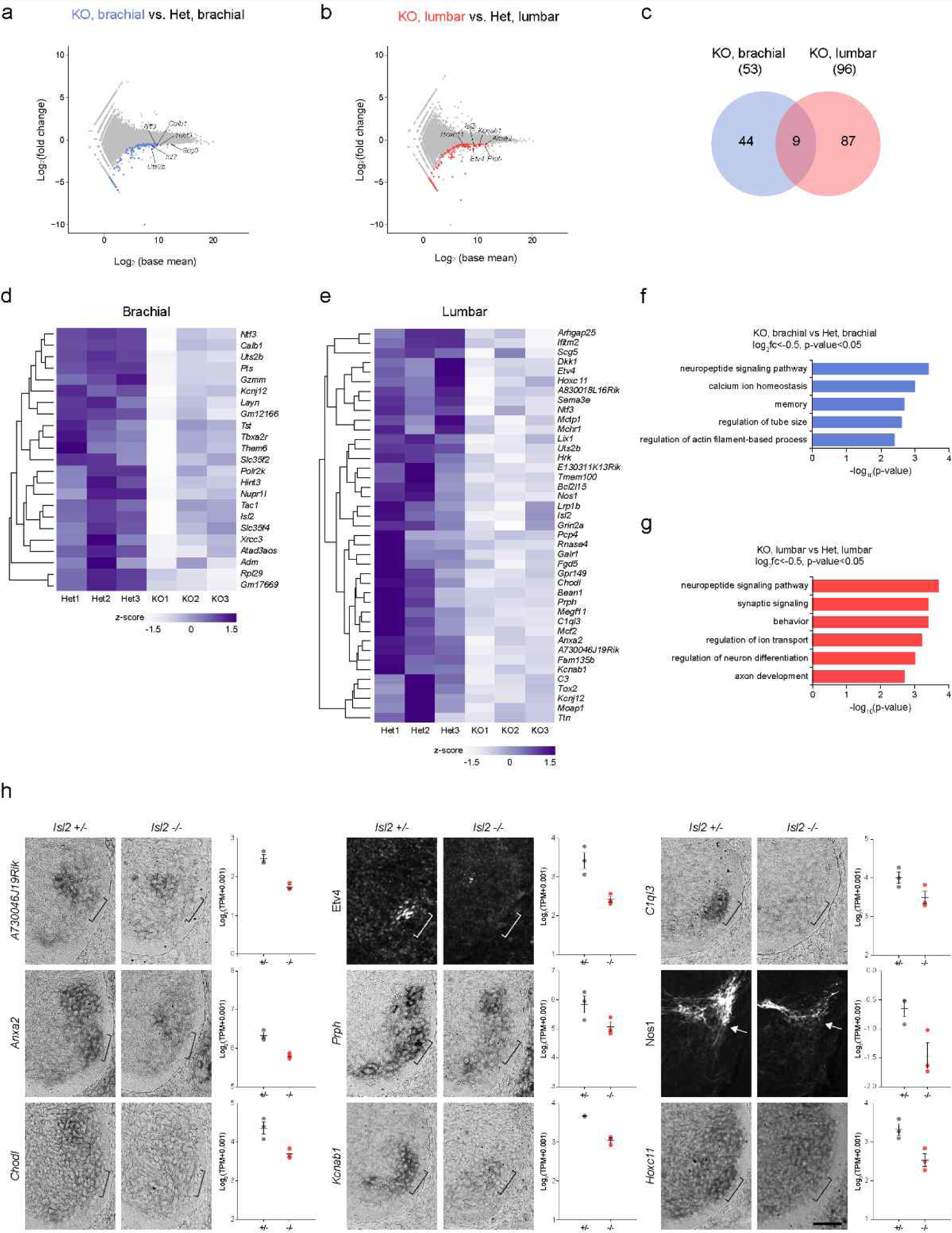
Transcriptome analysis of *Isl2*-deficient brachial and lumbar MNs at E12.5. (a, b) MA plots highlighting top DEGs among brachial and lumbar motor neurons in E12.5 *Isl2*-null and Isl2^+/−^ embryos. (c) Venn diagram of the downregulated genes of brachial and lumbar MNs in *Isl2*-null embryos. (d, e) Heatmaps of selected downregulated genes in the brachial and lumbar spinal cords of *Isl2*-deleted embryos. (f, g) Gene ontology (GO) analysis of DEGs downregulated at the brachial and lumbar levels in *Isl2* KO embryos. (h) *In situ* hybridization and immunohistochemistry of selected downregulated genes in E12.5 *Isl2*-deficient lumbar spinal cords and graphs of normalized read counts. Brackets indicate the ventrolateral motor pools, and arrows indicate PGC neurons. Data points of the graphs indicate normalized read counts from individual animals. SEM is shown. Scale bar, 50 μm.

To pinpoint the specific hindlimb motor pools affected by the downregulation of Isl2, we further analyzed the LMC motor pools using a published single-cell RNA-sequencing dataset from E12.5 mouse spinal cords (Amin et al., 2021). To analyze the motor pools in the rostral lumbar spinal cord (L1 to L4), we isolated the LMC subclusters that expressed high levels of *Hoxd9*, *Hoxa10*, *Hoxc10*, *Hoxd10*, *Hoxa11*, *Hoxc11*, and *Hoxd11* (Supplementary Figure 7). UMAP of 273 cells identified five clades: two LMCl and three LMCm subclusters (Supplementary Figure 7a–h). Subcluster LMCl.1 (sb.LMCl.1) expressed *Isl2* and *Lhx1* but not *Etv4* and was further divided into two subclusters: sb.LMCl.1.v, which consisted of V motor pools that expressed *Isl2*, *Lhx1*, and *Etv1*, and sb.LMCl.1.ta, which consisted of Ta motor pools that expressed *Isl2*, *Lhx1*, and *Nkx6-1*. Sb.LMCl.2 contained Rf, Tfl, and Gl motor pools that expressed *Lhx1*, *Nkx6-2*, and *Pea3*/*Etv4*. Likewise, sb.LMCm.1 was characterized by the expression of *Isl2*, *Isl1*, and *Etv1*, the molecular profiles of which were similar to the A/G motor pools. Subclusters sb.LMCm.2 and sb.LMCm.3 were both defined by the expression of *Isl2*, *Isl1*, and *Nkx6-1* and contained H and Gs motor pools, and sb.LMCm.2 (*Hoxd9*^high^ and *Aldh1a2*^high^) occupied a more rostral position than sb.LMCm.3 (*Hoxd9*^low^ and *Aldh1a2*^low^) (Supplementary Figure 7). Thus, we successfully defined five subclusters of LMC motor pools. Among the LMC subclusters, *Isl2* expression was higher in sb.LMC1.2 than in other subclusters, which suggested a close genetic relationship between *Isl2* and *Etv4* (Supplementary Figure 7b and d). Among the 87 DEGs selectively downregulated in *Isl2-*deficient lumbar LMCs, seven genes, *C1ql3*, *Etv4*, *A830018L16Rik*, *Barx2*, *Kcnab1*, *Hrk*, and *Prph*, overlapped with the enriched genes in subcluster sb.LMCl.2 (Supplementary Figure 7i). Therefore, the molecular signature of subcluster sb.LMCl.2, which included seven DEGs, could be transcriptionally regulated by *Isl2.* This may contribute to the establishment of the proximal motor pool.

### Defective arborization and sensorimotor connectivity of *Isl2-*deficient hindlimb motor pools

The downregulation of *Pea3* and the transcriptomic analysis of *Isl2-*deficient mice suggested that these mice have similar phenotypes to those found in *Pea3* mutant mice, such as scattered cell bodies, reduced dendrite patterning, and sensorimotor connectivity (Vrieseling & Arber, 2006). There were three motor pools that expressed Pea3 at the lumbar level of the spinal cord: Gl, Rf, and Tfl motor pools (Arber et al., 2000; De Marco Garcia & Jessell, 2008). Therefore, we examined the dendrite formation of *Isl2-*deficient motor pools by injecting rhodamine-dextran (Rh-Dex) retrograde tracer into individual muscles at P4. In the wild-type animals, the Gl motor pools exhibited a stereotyped elongated crescent shape of the dextran-labeled dendritic arbor at the ventrolateral position in L3–L4 spinal cords (Figure 6a). However, in *Isl2* mutant mice, dextran-labeled Gl motor pools split into two groups: 61% of cells were located at the correct ventrolateral position, and 39% were located at the ectopic ventromedial position. Regardless of position, *Isl2-*deficient Gl motor pools showed shorter and more randomly oriented dendritic arbors (Fig. 6a). We next determined the density of sensory synaptic contacts in control and *Isl2*-null mice. Rh-Dex was injected into the Gl muscle at P16, and the number and density of vesicular glutamate transporter 1 (vGluT1), a marker of proprioceptive synapses, were analyzed at P21. We observed a reduction in the number and density of vGluT1^+^ boutons on the soma of Rh-Dex^+^ Gl motor neurons in *Isl2*-null mice (Figure 6b). These results indicated that the Gl motor pools received fewer proprioceptive sensory input boutons in the absence of *Isl2*. Collectively, *Isl2* KO mice displayed reduced and misoriented dendrites of Gl motor pools with defective sensorimotor connections.

**Figure 6.**
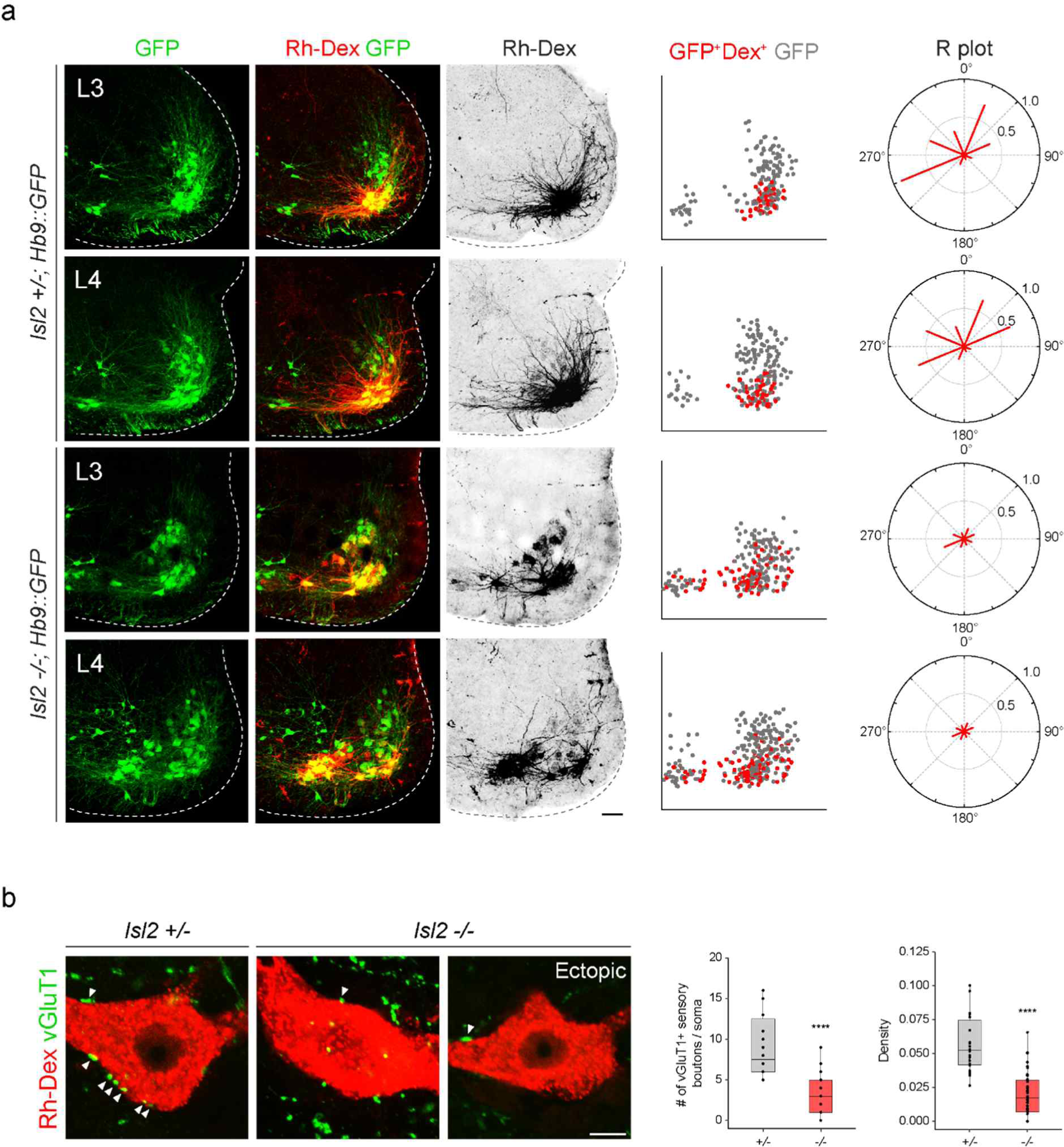
Impaired proprioceptive sensory nerve connectivity in the Gl motor pools of *Isl2* mutant mice. (a) Representative dendritic arbors of Gl motor pools retrogradely labeled with rhodamine-dextran (Rh-Dex). Spatial plots show soma position of Hb9::GFP^+^ (gray) and dextran-labeled Hb9::GFP^+^ motor neurons (red). Radial (R) plots show average dendritic membrane density per octant (red bars) from the motor somata (three animals per group; see Supplementary Table S2 for detailed sample sizes). Scale bar, 100 μm. (b) The contact of vGluT1^+^ Gl sensory boutons with Rh-Dex-labeled Gl motor neurons (arrowheads) in P21 animals. The number of sensory boutons per cell body (n = 14–23 cells from three animals per genotype; box plots showing feature distribution represent median (center line), first and third quartiles (box boundaries), and 10th and 90th percentiles (whisker); unpaired Student’s t-test; ; see Supplementary Table S2 for detailed n and statistics). \*\*\*\**p* < 0.0001. Scale bars, 10 μm.

### Aberrant neuromuscular junction (NMJ) formation and terminal axon branching in *Isl2*-null mice

To determine whether axon projections of mispositioned motor columns reach their target muscles in the limb, we traced LMC axon trajectories of *Isl2*-null mice carrying *Hb9::GFP*. LMC axons reach the base of the limb and diverge to either dorsal or ventral limb trajectories. The major brachial and lumbar plexus were largely normal in *Isl2*-null mice (Supplementary Figure 8a-c). However, axon outgrowth in response to glial cell-derived neurotrophic factor (GDNF) was significantly reduced in only lumbar motor axon explants of *Isl2*-null mice grown in vitro (Supplementary Figure 8d). We then examined the formation of the NMJ in the hindlimb muscles. Because NMJs mature during the first postnatal weeks, we examined the gluteus muscles at three different time points: at P0 when NMJ formation is ongoing, at P14 when synaptic elimination begins, and at 3 weeks when NMJs are monoinnervated and mature (Wyatt & Balice-Gordon, 2003). At P0, Hb9::GFP-labeled main motor axon bundles were comparable across *Isl2-*deficient Gl muscles; however, the branches originated from the main bundles, and the terminal arbors were drastically diminished. The number of NMJs was reduced to 84% in the *Isl2* mutant group, and the length and complexity of the secondary branches were significantly reduced (Figure 7a, c, and d). Higher magnification views revealed that numerous short aberrant sprouts and NMJs had developed near the primary branches in *Isl2-*deficient muscles, which was not observed in the control group (Figure 7b). The endplate band area occupied by motor axons was reduced to 55%, and the endplate band area occupied by AChR clusters marked by α-bungarotoxin (BTX) was reduced to 58% in the Gl muscles of the control group (Figure 7e and f). At P14, a drastic loss in AChR clusters and excessive growth of motor axons that bypassed the AChR clusters were observed in *Isl2*-null mice (Figure 7g–i). When the few surviving NMJs were visualized during the period of synaptic elimination, a significant portion of nerve terminals were aberrant in *Isl2*-null mice, showing faint AChR clusters, polyinnervation, denervation, and swelling of axons (Figure 7j–m, and o). The area of the NMJ defined by the nerve terminal and AChR cluster areas were also enlarged in the muscles of *Isl2*-null mice (Figure 7j, n). Fragmented and less compact AChR clusters were more abundant in *Isl2*-null mice than in heterozygote mice (Figure 7j, p, q). Other muscles innervated by non-Pea3-expressing motor pools had normal NMJ development (Supplementary Figure 9). Taken together, axon terminal arborization of the NMJ was selectively affected in Pea3-expressing motor pools when *Isl2* expression was compromised.

**Figure 7.**
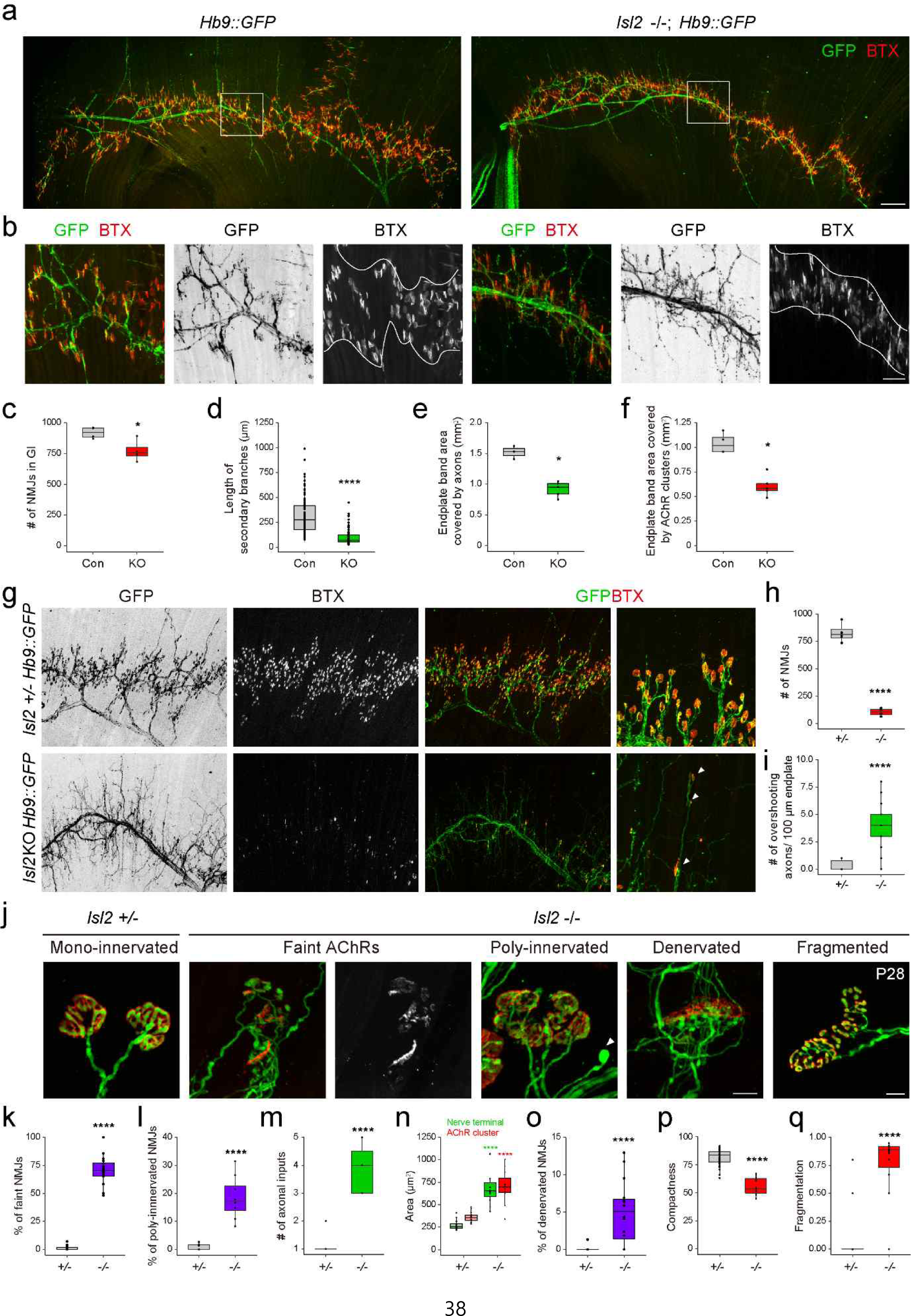
Reduced terminal motor axon branching in the gluteus (Gl) muscles of *Isl2* mutants. (a) Visualization of motor axons and neuromuscular junctions (NMJs) in P0 Gl muscles with Hb9::GFP (green) and α-bungarotoxin (BTX, red) immunoreactivity. (b) The boxed regions in (a) are shown at a higher magnification. The boundary of the region occupied by NMJs is marked by white lines. (c–f) The number of NMJs (c), secondary branch length (d), end plate area coverd by motor axons (e), and end plate area coverd by AChR clusters (f). (g) Images of motor axons and NMJs in P14 Gl muscles with Hb9::GFP (green) and α-bungarotoxin (BTX, red) immunoreactivity. Arrowheads indicate atrophic NMJs on overshooting axons. (h, i) The number of AChR clusters (h) and axons extending beyond AChR clusters at P14 (i). (j) Representative examples of various NMJs with morphological defects at a higher magnification at P14. For compactness and fragmentation analysis, P28 NMJs were analyzed. A swelling axon is indicated by an arrowhead. (k–q) Percentage of faint NMJs, polyinnervated NMJs, the number of axonal inputs, the area of AChR clusters and the nerve terminal, the percentage of denervated NMJs, compactness and fragmentation of AChR clusters were measured. n = 3 animals per group; box plots showing feature distribution represent median (center line), first and third quartiles (box boundaries), and 10th and 90th percentiles (whisker); unpaired Student’s t-test in (c, d, e, f, h, n); Mann-Whitney test in (i, k, l, m, o, p, q); see Supplementary Table S2 for detailed n and statistics. \**p* < 0.05, \*\*\*\**p* < 0.0001, n.s = not significant. Scale bars: (a) 200 μm, (b) 50 μm, (g) 200 μm (for low magnification images), 50 μm (for high magnification images), and (j) 10 μm.

### Impaired hindlimb movement in *Isl2-*deficient mice

A previous study reported that *Isl2*-null mice cannot survive after birth. However, we revealed that approximately 10% survived, albeit with lower body weight (Figure 8e, Supplementary Figure 10a, and Supplementary Table S1) (Thaler et al., 2004). Interestingly, newborn *Isl2* KO mice exhibited rigid hindlimbs that were parted widely, unlike their forelimbs, during free walking (Supplementary Movie 1). Adult *Isl2* KO animals also maintained an unnatural extended hindlimb posture when suspended by the tail and during walking (Figure 8a and Supplementary Movie 2). Footprint analysis showed that *Isl2* KO mice had abnormal gait of the hindlimb and broader width than that of the control group, while the gait of the forelimbs was normal (Figure 8b and c). A rigid and broad posture of the hindlimb may be a sign of proximal muscle weakness. Indeed, the muscle mass of the hip muscles, including the Gl and Rf muscles, was significantly reduced in *Isl2* KO animals (Figure 8d and e). X-ray imaging and whole skeleton staining with alizarin red and alcian blue showed that the overall bone structure and growth were normal in these animals (Supplementary Figure 10b and c). We then conducted electromyographic (EMG) analysis to measure hindlimb muscle activity in *Isl2* KO mice. Recordings from the Gl muscles in free-walking *Isl2* KO mice showed lower firing frequency, fewer single-motor-unit potentials, and shorter burst activity duration than the heterozygote animals (Figure 8f–j). Taken together, *Isl2* mutant mice displayed abnormal rigid hindlimb movement, which was accompanied by impaired EMG activity.

**Figure 8.**
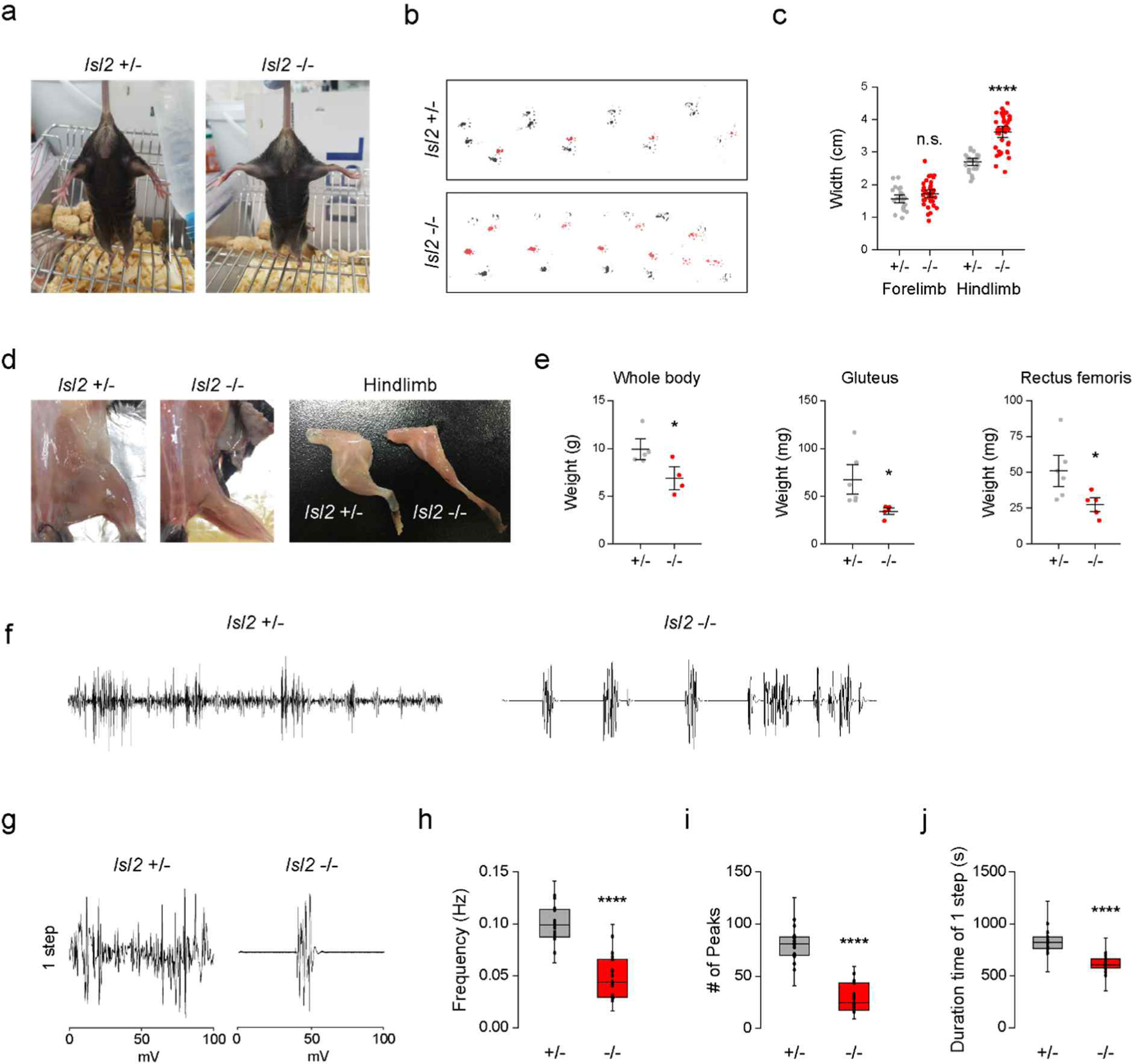
Defective hindlimb gait of *Isl2* mutants. (a) Representative images of the tail suspension test show a rigid wide-open posture of the hindlimbs in *Isl2* mutant mice. (b) The footprint patterns of 3-month-old *Isl2* +/− and *Isl2* KO mice. (c) Measurement of the stride width of forelimbs and hindlimbs (n = 4 and 8 animals per genotype; SEM is shown; unpaired Student’s t-test; see Supplementary Table S2 for detailed n and statistics). \*\*\*\**p* < 0.0001, n.s = not significant. (d) Representative gross appearance of lower limbs and dissected hindlimb samples of adult *Isl2* +/− and *Isl2* KO mice. (e) Measurement of body and muscle weight of the gluteus and rectus femoris muscles (n = 4 animals for heterozygote, n = 5 animals for KO; SEM is shown; unpaired Student’s t-test; see Supplementary Table S2 for detailed n and statistics). \**p* < 0.05. (f) Representative EMG recordings of the Gl muscles during free-walking. (g) Signature of individual footsteps. (h–j) Quantification of the average frequency, number of single-motor-unit potentials, and burst activity duration (three animals per group; box plots showing feature distribution represent median (center line), first and third quartiles (box boundaries), and 10th and 90th percentiles (whisker); unpaired Student’s t-test; see Supplementary Table S2 for detailed n and statistics). \*\*\*\**p* < 0.0001.

## Discussion

The acquisition of specialized motor pools and the specificity of target muscle connections with precision are key steps for effective motor control. We demonstrated that *Isl2* is critical for the clustering of motor pool subsets and the formation of dendritic arborization and NMJs in the hindlimb proximal muscles by demonstrating that *Isl2*-deficient mice display muscle rigidity and abnormal co-ordination in the limbs. We discuss our findings in the context of genetic programs of Isl2 that direct these connections in a motor pool-specific manner.

### Motor pool-specific contribution of Isl2 during motor neuron development

Isl2 is broadly expressed in motor neurons during motor neuronal development, similar to the panmotor neuronal expression of Isl1 but with a slight delay (Amin et al., 2021; Delile et al., 2019; Thaler et al., 2004). Isl1 is initially detected in all newborn motor neurons shortly after they exit the mitotic cycle at E9.5 (Pfaff et al., 1996; Thaler et al., 2004). However*, Isl2* begins its expression slightly later, at E10.5, in motor neurons (Thaler et al., 2004). Recent scRNA-seq and bulk-seq analysis studies have also revealed that *Isl2* expression arises in immature motor neurons, and expression is highest in the MMC and moderate in other motor columns (Amin et al., 2021; Catela et al., 2022; Delile et al., 2019; W. Wang et al., 2022). Therefore, the gradual increase in *Isl2* expression among immature motor neurons indicates functions that are distinct from those of Isl1. Our careful observation revealed that the clustering of motor neurons was disorganized in *Isl2* mutant spinal cords; MMC neurons were scattered at all segmental levels, and some LMCl motor pools were disorganized at the hindlimb level. In particular, Pea3-expressing LMCl motor pools lost their motor pool marker expression and displayed impaired NMJs. Notably, MMC and LMCl populations in a normal condition had relatively low or no expression of Isl1 but retained a substantial level of Isl2 expression, as shown by our in vivo results and scRNA-seq dataset. Thus, differential expression of Isl1 and Isl2 is critical for grouping motor neuronal subtypes, and the contribution of Isl2 is more substantial among populations in which either Isl2 expression is high, as in MMC neurons, or Isl1 expression is low, as in LMCl neurons.

By carefully dissecting the motor neuron clusters in published scRNA-seq data, we were able to successfully segregate the clades of LMC motor pools into six subclusters that encompassed most of the major motor pools at the rostral lumbar segment. Among these, *Isl2* expression was particularly prominent in sb.LMCl.2, which contained Gl, Tfl, and Rf motor pools. It was notable that sb.LMCl.2 cells commonly expressed *Pea3* and innervated the proximal hindlimb muscles. Moreover, Liau et al. described the molecular codes for ten subclusters for lumbar LMC MNs (Liau et al., 2021). Surprisingly, the LMCl subcluster called c/rl4 (*Lhx1*^+^ *Etv4*^+^), the molecular features for which matched our sb.LMCl.1, shared ten genes (*Anxa2*, *A730046J19Rik*, *Chodl*, *C1ql3*, *Etv4*, *Kcnab1*, *Lix1*, *Megf11*, *Prph*, and *Uts2b*) with those downregulated in the RNA-seq of *Isl2* KO embryos. Similarly, Wang et al. showed that LMC clusters express *Etv4*, which could be subdivided further (W. Wang et al., 2022). Thus, the contribution of *Isl2* appears to be substantial among motor pools innervating proximal hindlimb muscles.

### Isl2-dependent activation of the GDNF-Pea3 pathway in hindlimb LMC motor pools

The role of Pea3 in settling motor pool identity and sensorimotor connectivity has been well characterized, especially in brachial motor pools (Arber et al., 2000; Haase et al., 2002; Livet et al., 2002; Vrieseling & Arber, 2006). In this study, we observed that *Isl2*-null mice selectively lost *Pea3* expression in the lumbar spinal cord and thus displayed a phenotype almost identical to defects reported in *Pea3*-null mice, including mispositioned cell bodies, defective arborization of dendrites and NMJs, and reduced sensorimotor connectivity. Abolishment of the *Pea3* transcript was also observed in acute knockdown of *Isl2* in the chicken neural tube and in the spinal cords of motor neuron-specific *Isl2* cKO mice, which indicated that Isl2 positively regulates *Pea3* expression in a cell-autonomous manner. Notably, Pea3 expression in the CM motor pool at the brachial level was intact in *Isl2*-null mice and conditional mice. Perhaps Isl1 presents in CM motor pools, which belong to the LMCm, plays a redundant role that is similar to that of Isl2 in these populations. Indeed, the interchangeable role of the paralogs of LIM-HD transcription factors has been demonstrated in various cellular contexts. For instance, structural and biochemical studies have demonstrated that Lhx-Isl complexes show partial redundancy between Lhx3 and Lhx4, and between Isl1 and Isl2 (Gadd et al., 2011). Likewise, the coexpression of Lhx and Isl factors is equally effective in the ectopic generation of motor neurons in chicken embryos, and rescue experiment in zebrafish showed that two islet factors are equally potent (Hutchinson & Eisen, 2006; Kim et al., 2016; Song et al., 2009). Finally, genetic studies have revealed that the elimination of both *Lhx3* and *Lhx4*, but neither alone, abolishes motor neuron differentiation (H. Lee et al., 2015; Sharma et al., 1998). Thus, spatiotemporal dynamic expression among paralogs of various LIM-HD transcription factors enables segment-specific development of motor neurons.

Another important player that governs the development of Pea3-expressing motor pools is GDNF, a potent survival factor for motor neurons during the initial stage of motor neuron development(Henderson et al., 1994). GDNF expression subsequently becomes restricted in specific muscles, in which it acts as a peripheral signal to turn on *Pea3* expression in the motor pools innervating them (Haase et al., 2002). When GDNF signaling is disrupted, as in *GDNF* signaling mutants, motor pool positioning and innervation of target muscles are severely affected, which is similar to the defects exhibited by *Pea3* mutant mice (Haase et al., 2002; Livet et al., 2002). In addition, the elimination of the GDNF receptor in conditional *Ret* KO mice results in an abnormal hindlimb position with a club-foot phenotype because of abnormal axon innervation of motor neurons to the hindlimb, which emphasizes the importance of GDNF signaling in the control of hindlimb movement (Kramer et al., 2006). Consistent with these findings, we demonstrated that *Isl2-*deficient motor axons failed to grow in response to GDNF in vitro and lacked Pea3 expression, which indicated that Isl2 is involved in the GDNF-Pea3 pathway. When either gene is absent, the phenotypes of hindlimb motor pools were almost identical: scattered motor pools, disorganized and misoriented dendritic fields, and sensorimotor connectivity with inappropriate targets. Thus, Isl2 gates GDNF signaling to specific hindlimb motor pools by inducing Pea3 expression.

### Transcriptomic analysis to uncover Isl2-dependent genes for motor pool formation

Our transcriptomic analysis and in vivo validation of candidate targets revealed that numerous genes involved in motor neuronal differentiation and diseases were indeed under the control of Isl2. Among the 96 DEGs downregulated in *Isl2* mutant spinal cords, 31 genes (i.e., *Adra2c, Arhgap25, C1ql3, C3, Cck, Chrm5, Dkk1, Etv4, Fstl3, Gabra5, Gpr149, Grin2a, Hrk, Htr4, Kcnab1, Kcnj12, Mcf2, Mchr1, Mctp1, Nos1, Ntf3, Oprd1, P2rx4, Pcp4, Pnmt, Prph, Pvalb, Scnn1a, Sema3e, Tmem100, Tmsb15b2,* and *Tnc*) were associated with axon- or synapse-related GO terms, or their protein expression was found in axons or synapses (Dhaese et al., 2009; Fink, Lopez-Giraldez, Kim, Strittmatter, & Cafferty, 2017; Martinelli et al., 2016; Molinard-Chenu et al., 2020; T. Wu et al., 2019). In addition, 14 genes (i.e., *Anxa2, Bcl2l15, C3, Chodl, Fam135b, Hrk, Lix1, Lrp1b, Moap1, Pcp4, Prph, Rnase4, Scg5,* and *Scnn1a*) were reported as ALS- or motor neuron disease-associated genes in literatures (Benoit et al., 2013; Ghavami et al., 2014; Gros-Louis et al., 2004; Lederer, Torrisi, Pantelidou, Santama, & Cavallaro, 2007; Li et al., 2013; Orr et al., 2020; Sasongko et al., 2010; Sheila et al., 2019; Shu, Wei, & Zhang, 2022; Sleigh et al., 2014; Su, Wang, & Meng, 2022; Wei, Blundon, Rong, Zakharenko, & Morgan, 2011; T. Wu et al., 2019). Therefore, approximately 40% of *Isl2-*dependent DEGs were involved in the maturation and pathological processes of motor neurons. *Prph*, a neuronal intermediate filament protein implicated in ALS, showed the largest fold-changes (<−0.5 log_2_-fold change) in our transcriptomic analysis and was selectively downregulated in specific motor pools in *Isl2* mutant mice. In addition, earlier extinction of *Prph* in developing motor neurons correlated with progressive degeneration of spinal motor neurons and NMJs, which indicated that the defects observed in *Isl2* mutant mice are relevant to early-onset progressive motor neuron disorders such as SMA and ALS (Amin et al., 2015; Amin et al., 2021; Beaulieu & Julien, 2003; Chen, Sayana, Zhang, & Le, 2013; Gros-Louis et al., 2004; Sabbatini et al., 2021). Similarly, C-type lectin chondrolectin (*Chodl*) is a known marker for fast MNs and is involved in cell survival and neurite outgrowth in MNs (Enjin et al., 2010; Sleigh et al., 2014). *Chodl* is dysregulated in spinal muscular atrophy models, and KD of *Chodl* elicits stalling of motor axonal growth and reduction of muscle innervation (Zhong et al., 2012). C1ql3, a complement component 1, q subcomponent-like protein, has been shown to promote synapse formation and maintenance in certain excitatory neurons, and its absence results in fewer excitatory synapses with diverse behavioral defects (Martinelli et al., 2016). Given that these motor neuron disease-related genes were all selectively downregulated in specific motor pools that showed deteriorating features in the absence of *Isl2*, it is likely that proper regulation of these genes by Isl2 ensures proper motor neuron development, whereas dysregulation leads to motor neuron degeneration.

In summary, our findings suggest that pool-specific Isl2 activity is important for the fidelity of motor pool organization and motor circuit connectivity for hindlimb locomotion. Furthermore, Isl2 is responsible for the transcriptional control of a variety of genes involved in motor neuron differentiation, axon development, and synaptic organization, and its absence may give rise to a pathological condition of motor neurons.

## Materials and methods

### Mice

The *Isl2*-null and *Hb9::GFP* mice used have been described previously (Thaler et al., 2004). Wild-type C56BL/6 and CD-1 mice (6–8 weeks old) were purchased from Damul Science Co. (Daejon, Korea). To selectively delete *Isl2* from motor neurons, we generated a novel *Isl2* flox mouse (Cyagen, China). The flox strategy was designed by Cyagen using the following procedure. The *Isl2* gene (NCBI reference sequence: NM_027397; Ensemble: ENSMUSG00000032318) was located on chromosome 9 in mice. Exons 2–4 were selected as cKO regions. In the targeting vector, the Neo cassette was flanked by SDA sites, and DTA was used for negative selection. C57BL/6N ES cells were used for gene targeting. The KO allele was obtained after specific Cre-mediated recombination. Genotyping assays were designed by Cyagen. First-generation mice were heterozygous for *Isl2* flox expression. Heterozygous mice were bred to generate mice with homozygous *Isl2* flox expression. The *Isl2* flox mouse line was maintained as a homozygous line for breeding with Cre strains. For the Cre line, a motor neuron-specific Olig2-Cre line was used (Dessaud et al., 2007; Kong et al., 2015). To obtain *Isl2^f/KO^*; Olig2-Cre mice, male *Isl2^+/−^*; Olig2-Cre mice were crossed with female *Isl2^f/f^* mice. The *Isl2^flox/KO^* or *Isl2^flox/+^*; Olig2-Cre mice from the same litters were used as normal controls. All experiments used protocols approved by the Animal Care and Ethics Committees of the Gwangju Institute of Science and Technology (GIST). The day when a vaginal plug was detected was designated embryonic day 0.5 (E0.5).

### Immunohistochemistry and *in situ* hybridization

Immunohistochemistry and *in situ* hybridization were performed as described previously(Song et al., 2009). The following antibodies were used: rabbit and guinea pig anti-HB9 (Harrison, Thaler, Pfaff, Gu, & Kehrl, 1999), rabbit anti-GFP (Invitrogen), mouse anti-GFP (Sigma), rabbit anti-Isl1/2 (Pfaff et al., 1996), mouse anti-Isl2 (Santa Cruz), guinea pig anti-Lhx3 (Thaler et al., 2004), goat anti-ChAT (Chemicon), rabbit anti-Foxp1 (Abcam), goat anti-Nkx6.1 (R&D Systems), goat anti-Isl1 (R&D Systems), rabbit anti-Lhx1 (Abcam), guinea pig anti-Scip (Dasen et al., 2005), rabbit anti-Pea3 (Dasen et al., 2005), rabbit anti-Tetramethylrhodamine (Invitrogen), guinea pig anti-vGluT1 (Millipore), rabbit anti-nNOS (Diasorin). For neuromuscular junction staining, muscles were harvested and were prepared for immunostaining as whole-mount samples or transverse sections. Samples were immunostained with rabbit anti-GFP (Abcam) and Alexa Fluor 555 *α*-BTX (Invitrogen). For *in situ* hybridization, embryonic mouse cDNA at E12.5 was used to generate riboprobes for *Aldh1a2, Er81*, *Prph*, *Anxa2*, *Chodl*, *A730046J19Rik*, *Kcnab1*, *C1ql3*, and *Hoxc11*, and HH stage 25 chicken embryonic cDNA was used to generate riboprobes for chicken *Raldh2* and *Pea3* using a one-step RT-PCR kit (Solgent).

### Chick in ovo electroporation

In ovo electroporation was performed as described previously (Kim et al., 2016). A DNA solution was injected into the lumen of the spinal cord of HH stages 10–12 chicken embryos, and the embryos were harvested at HH stage 25. To knock down *Isl2*, *Isl2* siRNAs (siRNA-461 sense 5’-GGA CGG UGC UGAACG AGAA-3’, siRNA-461 antisense 5’-UUC UCG UUC AGC ACC GUC C-3’; siRNA-605 sense 5’-GCU GCA AGG ACA AGA A-3’, siRNA-605 antisense 5’-UUC UUC UUG UCC UUG CAG C-3’) were electroporated. Harvested chicken embryos were processed for immunohistochemistry and *in situ* hybridization. The following antibodies were used: rabbit anti-GFP (Invitrogen), mouse anti-Isl2 (DSHB), and mouse anti-MNR2 (DSHB). Riboprobes for chicken *ALDH1A2* and *PEA3* were amplified from total RNA extracted from HH 25 chicken embryos using a one-step RT-PCR kit (Solgent). For image analysis, consecutive sections at the lumbar level were collected from each embryo. In each ventral quadrant, the number of motor neurons was counted, and the signal intensity was measured using the ImageJ software (NIH). The intensity values of the electroporated side were normalized to the value of the nonelectroporated side.

### Intramuscular injections of tracers

To label motor neuron dendrites, the target muscles were injected with Rh-Dex (3000 MW, Invitrogen). In brief, P4 mice were anesthetized on ice, and the muscles were exposed by making a small incision in the skin. Approximately 2 μl of 10% Rh-Dex was injected at a single site, and the skin was sutured. After 3 days, the spinal cords were analyzed for dextran labeling, and the muscles were isolated to check the injection site. To trace dendritic arborization patterns of the Gl motor pools, analyses were focused primarily on the L3 and L4 spinal segmental levels. Free-floating 80 μm-thick transverse sections were immunostained for further analysis.

For sensory bouton analysis, P16 mice were anesthetized, and the skin was incised to expose the muscle of interest. Muscles were injected at multiple sites with 10% tetramethylrhodamine dextran (3000 MW, Invitrogen) using a pulled glass microelectrode. At P21, the spinal cords were harvested and processed for immunostaining.

### EMG analysis

The electrodes were made from multistranded, Teflon-coated, annealed stainless steel wire (A-M systems, Cat. No. 793200) and soldered to an IC socket. After mice were anesthetized with avertin, electrodes were implanted subcutaneously into the middle of the left and right Gl muscles, and the connector was secured to the head. The mice were left in their cages for recovery for at least 2 days before recording began. After 2 days of recovery, the animal was placed in an open field where it could move freely. During locomotor activity, EMG signals were digitized and stored using the data acquisition software Clampex 9.2, and video recordings were acquired to monitor walking movement for 30 minutes. During the recording sessions, EMG and video data were recorded simultaneously. The EMG data were bandpass filtered at 200 Hz to 1 kHz using Clampfit 10.3. For EMG signal analysis, the number, frequency, and duration of EMG peaks over 1 second were measured and analyzed.

### Tail suspension test and gait analysis

For the tail suspension test, each mouse was suspended above the cage by its tail, and its posture was captured on a camera. For the gait analysis, 3-month-old *Isl2* KO and littermate female controls were used. After nontoxic water-based paints (forepaws in red and hindpaws in black) were applied to the mouse paws, the mice were allowed to walk on white paper. Three consecutive trials were performed. The stride widths of the forelimbs and hindlimbs were measured.

### RT-PCR

E12.5 embryonic spinal cords were harvested in ice-cold PBS. Total RNA was extracted using NucleoSpin RNA XS (MACHEREY-NAGEL). We used 10 ng of total RNA per 15 μl reaction for the RT-PCR reaction with each assay primer (*Gapdh*, 5’-GGA GAA ACC TGC CAA GTA TGA-3’, 5’-CCT GTT GCT GTA GCC GTA TT-3’, *Pea3* 5’-GGT GAT GGA GTG ATG GGT TAT G-3’, 5’GCC TGT CCA AGC AAT GAA ATG-3’). The PCR products were amplified using a one-step RT-PCR kit (Solgent). Band densitometry was quantified for *Pea3* and *Gapdh* using the ImageJ software.

### Data analysis

#### Image acquisition

For data analysis, images were captured using a Zeiss epifluorescent (LSM780 [Zeiss] and FV3000RS [Olympus]) confocal microscopes using the Axiovision and ZEN software (Zeiss) and the FV31S software (Olympus).

#### Radial plot

To analyze dendritic radiality, motor pools were subdivided into eight divisions, with the center aligned with the center of the motor pool. For each octant, the mean pixel intensity of dendrites, excluding the cell bodies, was quantitated using the ImageJ software and normalized to the octant with the highest value.

#### Contour density plot

Co-ordinates of the motor neuron soma were acquired with respect to the midline using the ImageJ software. Variations in co-ordinate values from different spinal cord sections were normalized according to the size and shape of the spinal cord. Distribution contours were plotted using the “ggplot2” package in R-4.0.5.

### Quantification of sensory boutons

For quantitative analysis of vGluT1^+^ sensory boutons in Rh-Dex^+^ Gl motor pools, spinal cords were sectioned at a thickness of 40 μm. A z-series of 0.74 μm optically scanned confocal images using a 30× objective were acquired for quantitative analysis. The number of vGluT1^+^ puncta within each somatic compartment was quantified across the Rh-Dex^+^ motor neurons. Motor neuron surface area was determined using the Image J software, and vGluT1^+^ synapses were counted manually (Y. Wang et al., 2019). The density of synaptic puncta was quantified by dividing the number of vGluT1^+^ synapses in the somatic compartment of Rh-Dex^+^ Gl motor pools by the length of each somatic compartment (Fletcher et al., 2017; Mentis et al., 2011; Vukojicic et al., 2019).

### Quantification of NMJ morphology and imaging

NMJ images were acquired on LSM780 (Zeiss) and FV3000RS (Olympus) confocal microscopes. To analyze NMJs, maximum intensity projections were created using the ZEN (Zeiss) and FV31S (Olympus) software. Low magnification images were acquired using 10× and 20× objectives. To assess NMJ morphology, confocal settings were optimized: 640 × 640 frame size, 30× objective, ×6.0 zoom, and 0.74 μm z-stack interval for quantification in P14 muscles; 800 × 800 frame size, 30× objective, ×4.5 zoom, and 0.74 μm z-stack interval for quantification in P28 muscles. The number of NMJs was quantified along the entire gluteal nerve within the gluteus muscles in P0 and P14. Quantification of the length of secondary branches, the end plate area occupied by motor axons or AChR clusters in P0 and the nerve terminal area, AChR cluster area, compactness and fragmentation in P14 and P28 was performed using the ImageJ software (Jones et al., 2016). The number of axons extending the border of the target muscles was calculated by quantifying the overshooting axons per 100 μm of endplate. The proportion of polyinnervated, faint, denervated NMJs was quantified within 3–4 random fields per animal. The proportion of polyinnervated NMJs was quantified in gluteal nerves adjacent to Tfl muscles. The number of axonal inputs was quantified in individual NMJs in P14.

### RNA-Seq and bioinformatics analysis

For the RNA-seq analysis using the ventral brachial and lumbar spinal cords, E12.5 embryonic spinal cords (three heterozygote animals and three homozygote animals) were harvested and trimmed around the *Hb9::GFP* reporter in ice-cold RNase-free PBS. Total RNA was extracted using NucleoSpin RNA XS (MACHEREY-NAGEL). RNA quality and quantity were measured using a 2100 Bioanalyzer System (Agilent). All samples acquired high-quality scores between 9.6 and 10 RIN. For library preparation, 500 ng of total RNA per sample was used to generate cDNA, which was amplified using the TruSeq Stranded Total RNA LT Sample Prep Kit (Gold). Sequencing was performed on the NovaSeq system. The processed reads were mapped to the mm10 mouse genome assembly using HISAT2 for further processing. Transcript assembly was performed using StringTie, and expression profiles of each sample were acquired. Analysis of DEGs was performed using DESeq2.

### Gene ontology (GO) analysis

GO analysis based on bulk RNA-seq data was performed using Metascape (http://metascape.org) (Zhou et al., 2019).

### Statistial representation in figures

Error bars represent SEM. Box plots show 25th percentile, median, and 75th percentile values, whiskers showing 10th and 90th percentile values, and symbols showing outlying values.

## Acknowledgments

This work was supported by grants from NRF (NRF-2018R1A5A1024261, NRF-2020R1A2C1006758, NRF-2022M3E5E8081194), KHIDI HI14C3484, and the GIST Research Institute in 2022. We thank Jawoon Yi, Jung Eun Kim, and Won-Young Lee for experimental setup support, and Myungin Baek for their critical comments. The pBS mouse ER81 [TJ#197] was a gift from Thomas Jessell (Addgene plasmid #16282; http://n2t.net/addgene:16282; RRID: Addgene_16282).

## Supplementary information

- Supplementary figure 1-10
- Supplementary table S1-S3
- Supplementary methods
- Supplementary video 1, 2

---

**Supplementary Figure 1.**
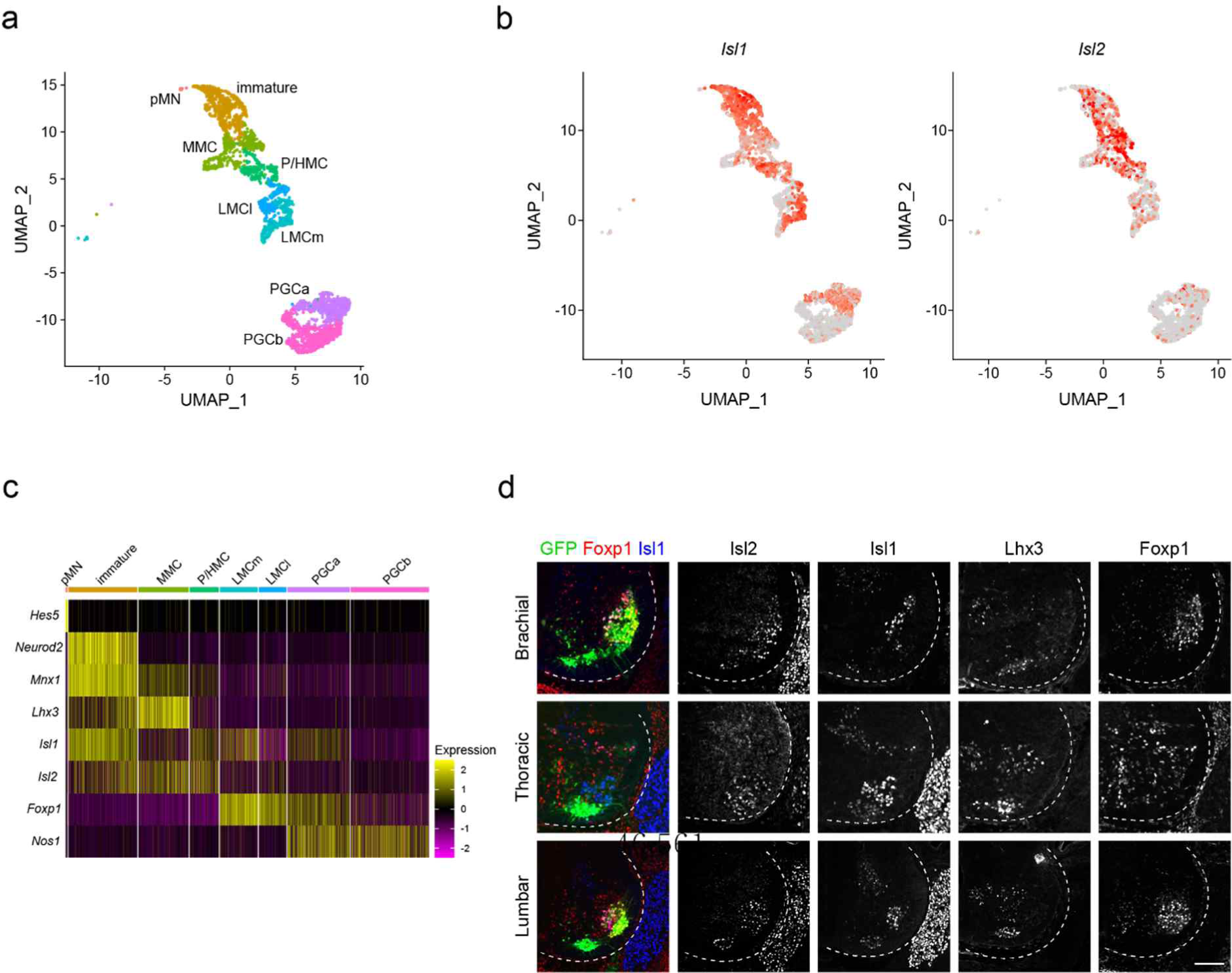
Spatiotemporal dynamics of *Isl2* expression and heterogeneity of motor neurons. (a) UMAP plot of scRNA-seq data from Amin et al., with cells colored according to clusters, as defined by Amin et al., showing MN-lineages, including pMN progenitors, immature motor neurons, MMC, P/HMC, LMCm, LMCl, PGCa, and PGCb. (b) UMAP plot showing differential expression of *Isl1* and *Isl2*. (c) Heatmap shows dynamic gene expression of *Isl1*, *Isl2*, and marker genes in motor neuron clusters. (d) Expression of Isl1, Isl2, Lhx3, Foxp1, and Hb9::GFP in E12.5 brachial, thoracic, and lumbar spinal cords. Scale bar, 100 μm.

**Supplementary Figure 2.**
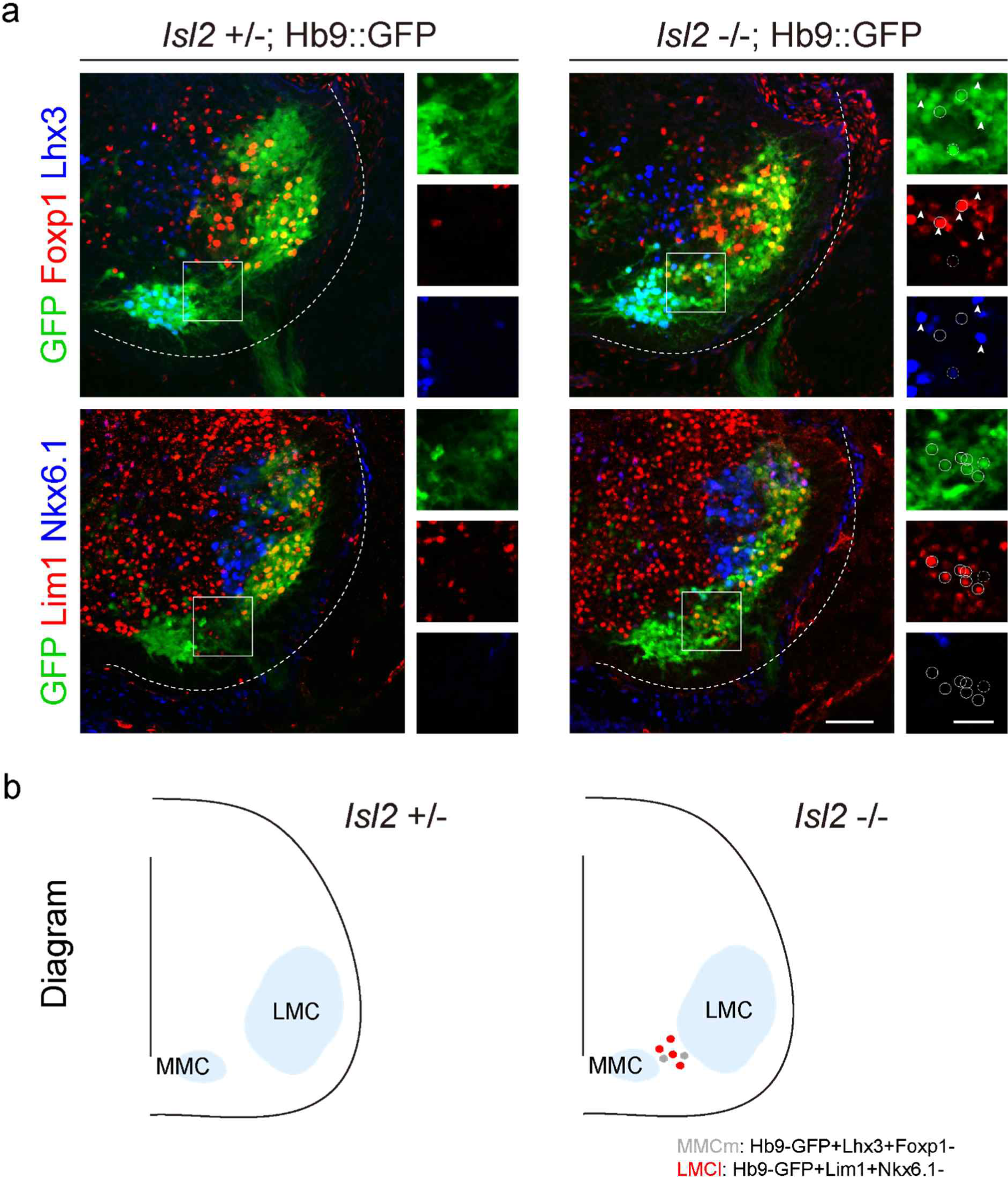
Characterization of ectopic motor neurons in *Isl2* knockout mice. (a) Representative images of ectopic motor neurons immunolabeled with various motor neuronal markers. High magnification images show MMC (arrowheads), LMCl (white circles), and unidentified (white dotted circles) cells in the region of ectopic motor neurons (n = 12 random fields, three animals per genotype). Scale bars, 50 μm (for low magnification images) and 25 μm (for high magnification images). (b) Summary diagram of ectopic motor neurons in *Isl2*-deficient mice.

**Supplementary Figure 3.**
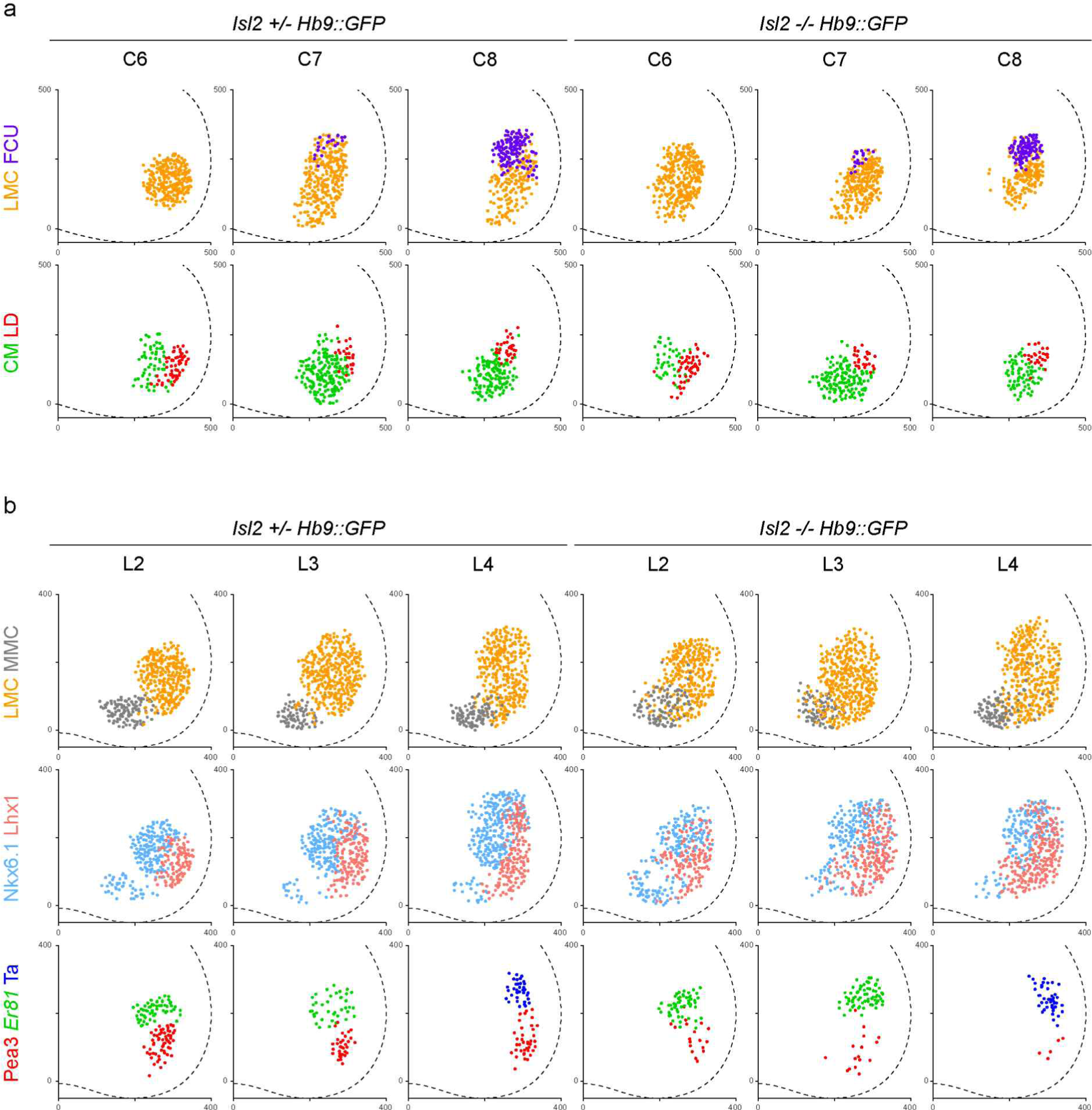
Spatial plots depict position of individual neurons comprising motor columns and motor pools indicated from three different animals shown in Figure 3.

**Supplementary Figure 4.**
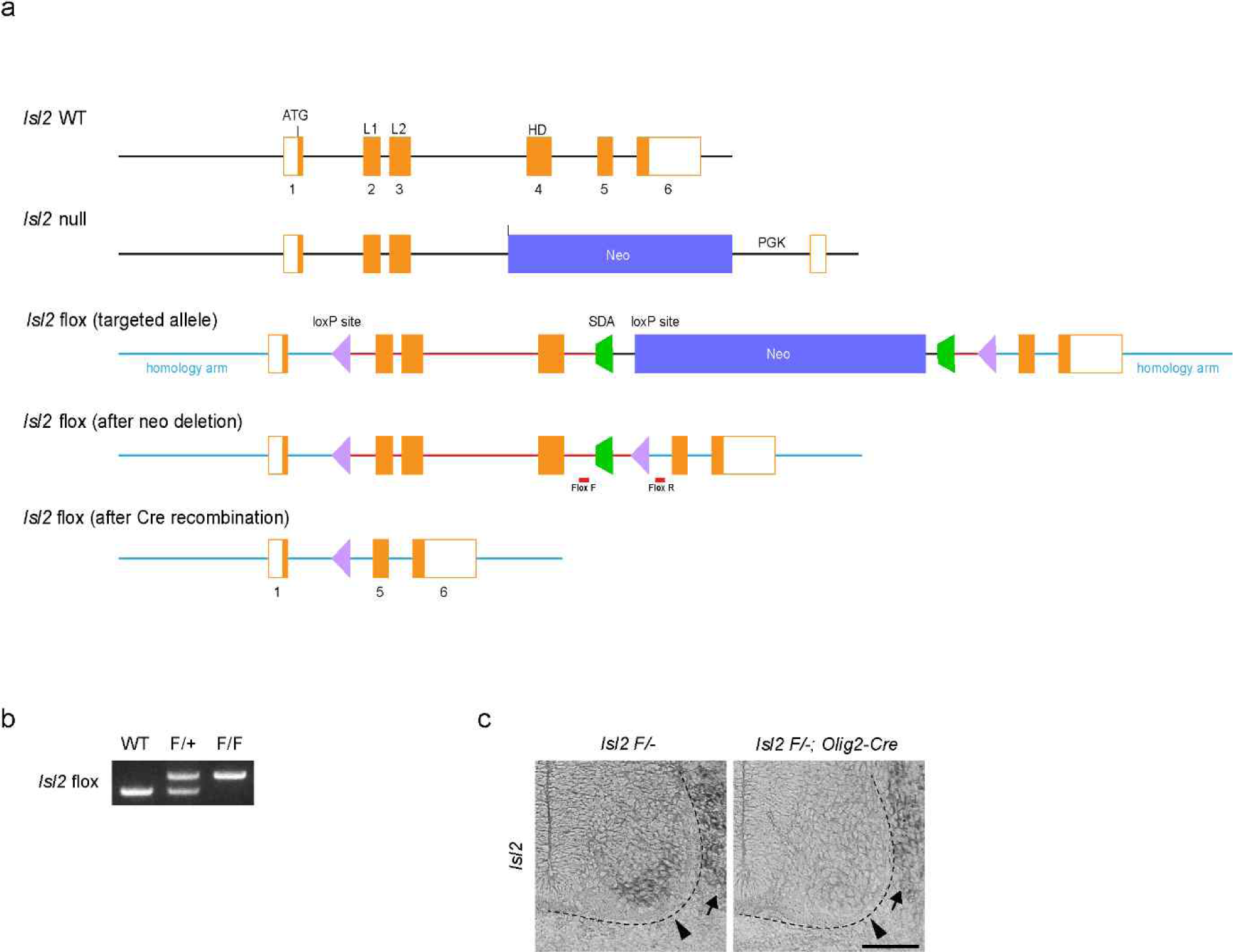
Generation of novel *Isl2* flox mouse line. *Isl2* flox mice were generated by Cyagen Biosciences (Guangzhou), Inc. (a) Gene targeting strategy used to generate *Isl2* flox mice. The flox strategy was designed to flank exon 2 to exon 4 with loxP sites. The knockout strategy is displayed for comparison, replacing three exons with the *neomycin* cassette (Thaler et al., 2004). Orange boxes, purple triangles, and green trapezoids represent exons with the exon number indicated, *loxP* sites, and the self-deletion anchor (SDA) sites, respectively. Blue lines denote the 5’ and 3’ homology arms approximately 2,722 and 2,933 kb long, respectively. The red line denotes the cKO region. Red lines show the location of the primers for genotyping. The F_1_ *Isl2^flox-neo/+^* mice were crossed with deleter mice to remove the *Neo* cassette in the germline. (b) Genotyping PCR confirmed the generation of wild-type heterozygous and homozygous pups. (c) *In situ* hybridization for *Isl2* mRNA in *Isl2 F/-* and *Isl2 F/-; Olig2-Cre* embryos at E12.5. Isl2. Note that mRNA expression of *Isl2* is diminished in motor columns (arrowheads) not in dorsal root ganglia (arrows). Scale bar, 100 μm.

**Supplementary Figure 5.**
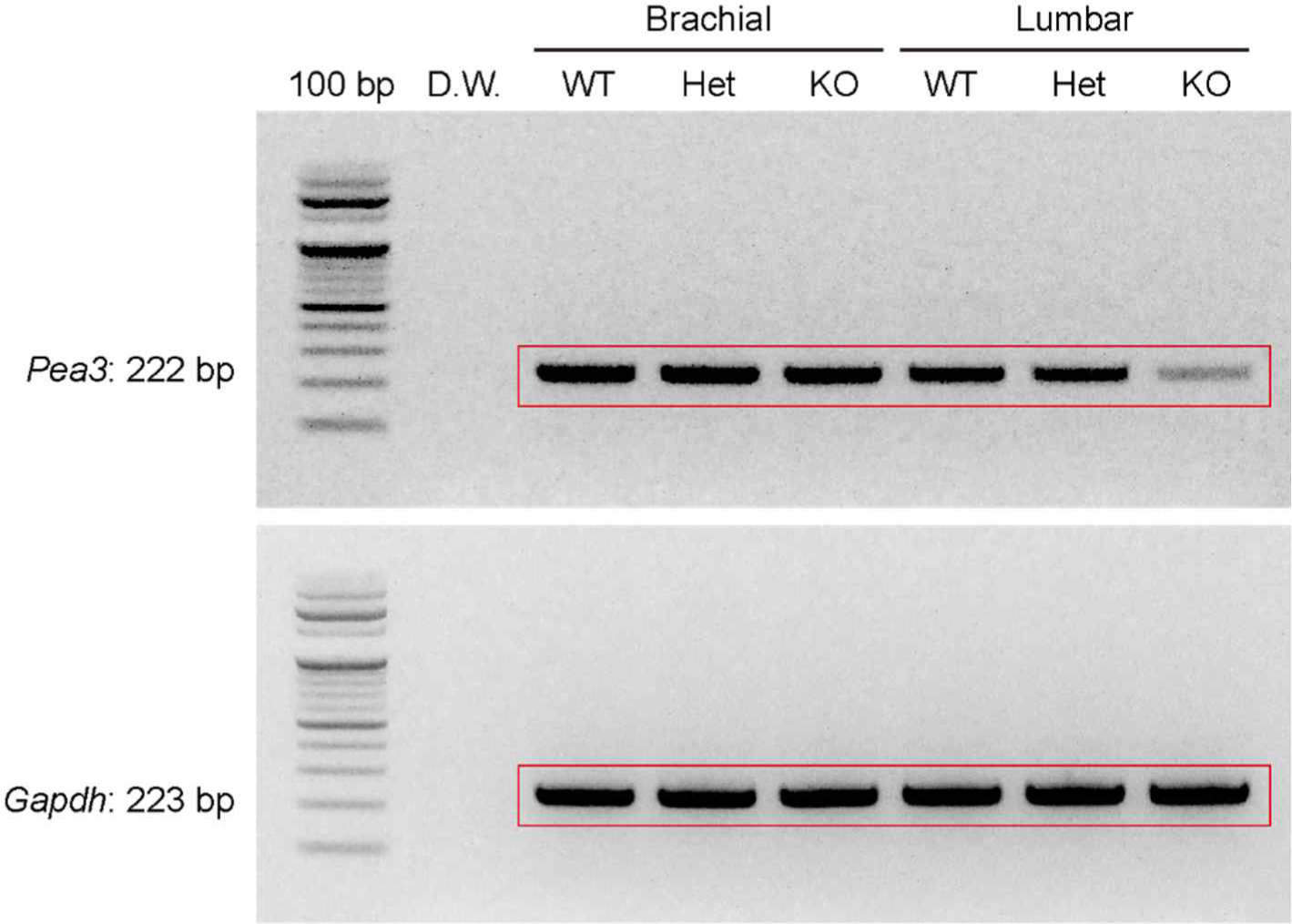
Original unprocessed images of RT-PCR results in Figure 4b. Original unprocessed images of *Gapdh* and *Pea3* RT-PCR in the brachial and lumbar spinal cords of E12.5 wild-type, *Isl2^+/−^*, and *Isl2*-null embryos.

**Supplementary Figure 6.**
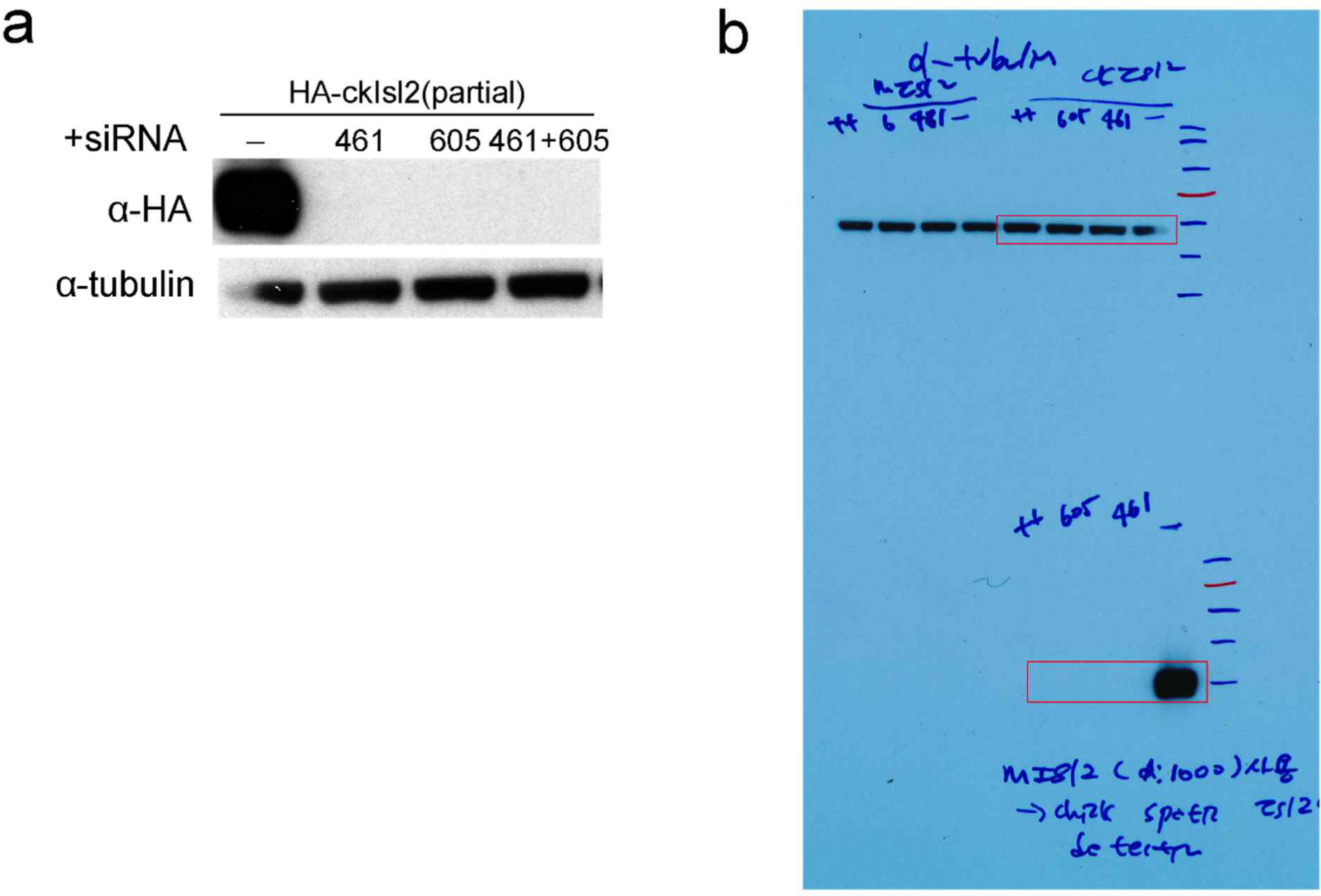
Knockdown (KD) of *Isl2* resulted in mislocalized motor neurons and lower *Pea3* expression. (a) KD efficiency of siRNA in HEK293T cells transiently transfected with N-terminal truncated chick *Isl2* (aa 38-356) with HA tag and *ckIsl2* siRNA 461 or 605, as indicated. Both siRNAs efficiently downregulated the protein expression of ckIsl2 determined by anti-HA immunoblot and siRNA 461 was used in Figure 4.

**Supplementary Figure 7.**
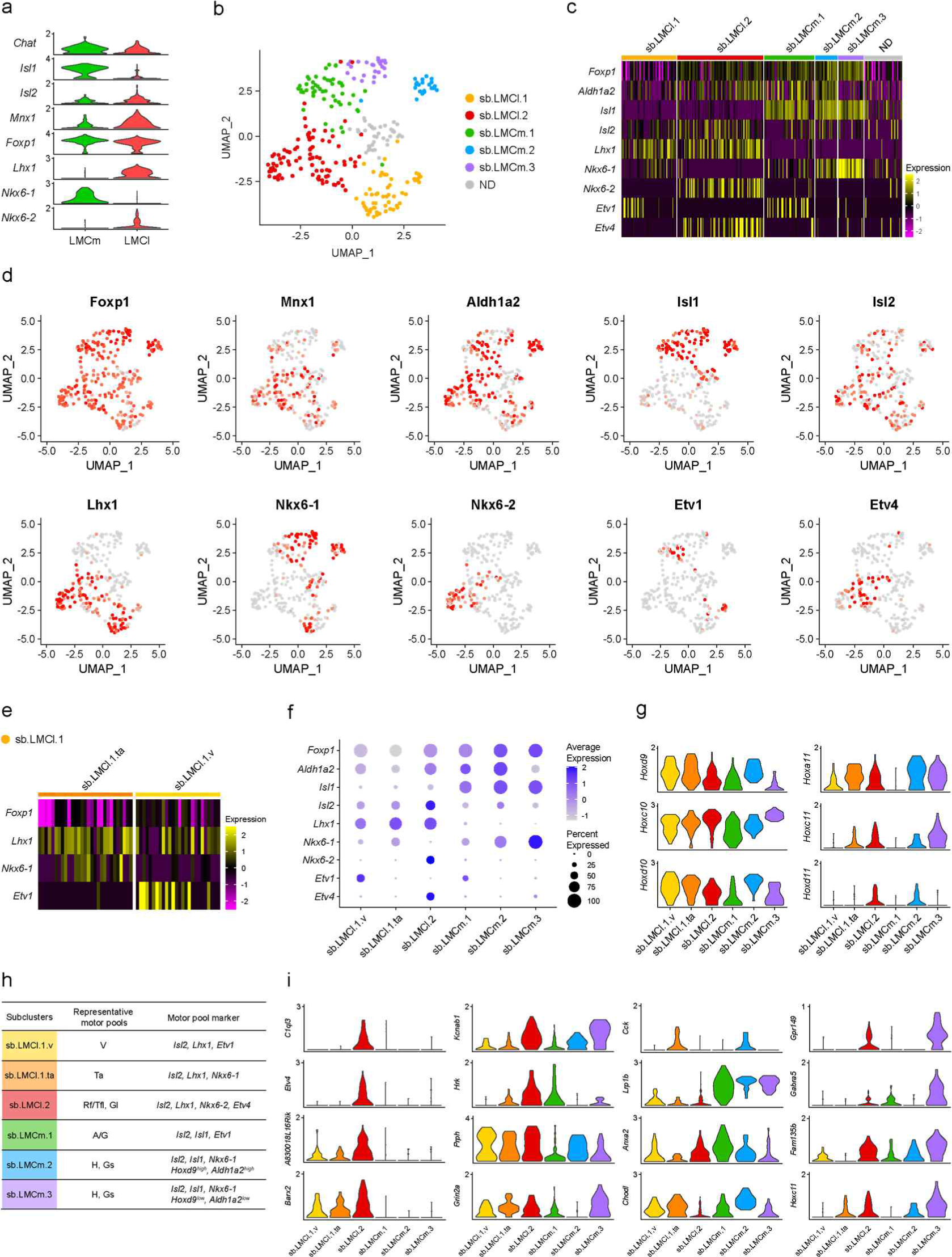
Identification of LMC clusters in which the gene signatures relate to *Isl2*. (a) Violin plots of MN subtype markers (*Chat*, *Isl1*, *Isl2*, *Mnx1*, *Foxp1*, *Lhx1*, *Nkx6-1*, and *Nkx6-2*) in LMCm and LMCl clusters. ND, not defined. (b) UMAP plot of cells from lumbar LMC clusters. (c) Heatmap showing marker genes for motor neuron subtypes and motor pools of six clusters. (d) UMAP plots showing gene expression of LMC and motor pool markers. (e) Heatmap showing marker genes for sb.LMCl.1.ta and sb.LMCl.1.v. (f) Dot plot showing expression patterns of MN markers of LMC subclusters. (g) Violin plots of *Hox* genes in LMC subclusters. (h) Summary of markers of LMC subclusters. (i) Violin plots of selected DEGs downregulated in *Isl2*-deficient spinal cords.

**Supplementary Figure 8.**
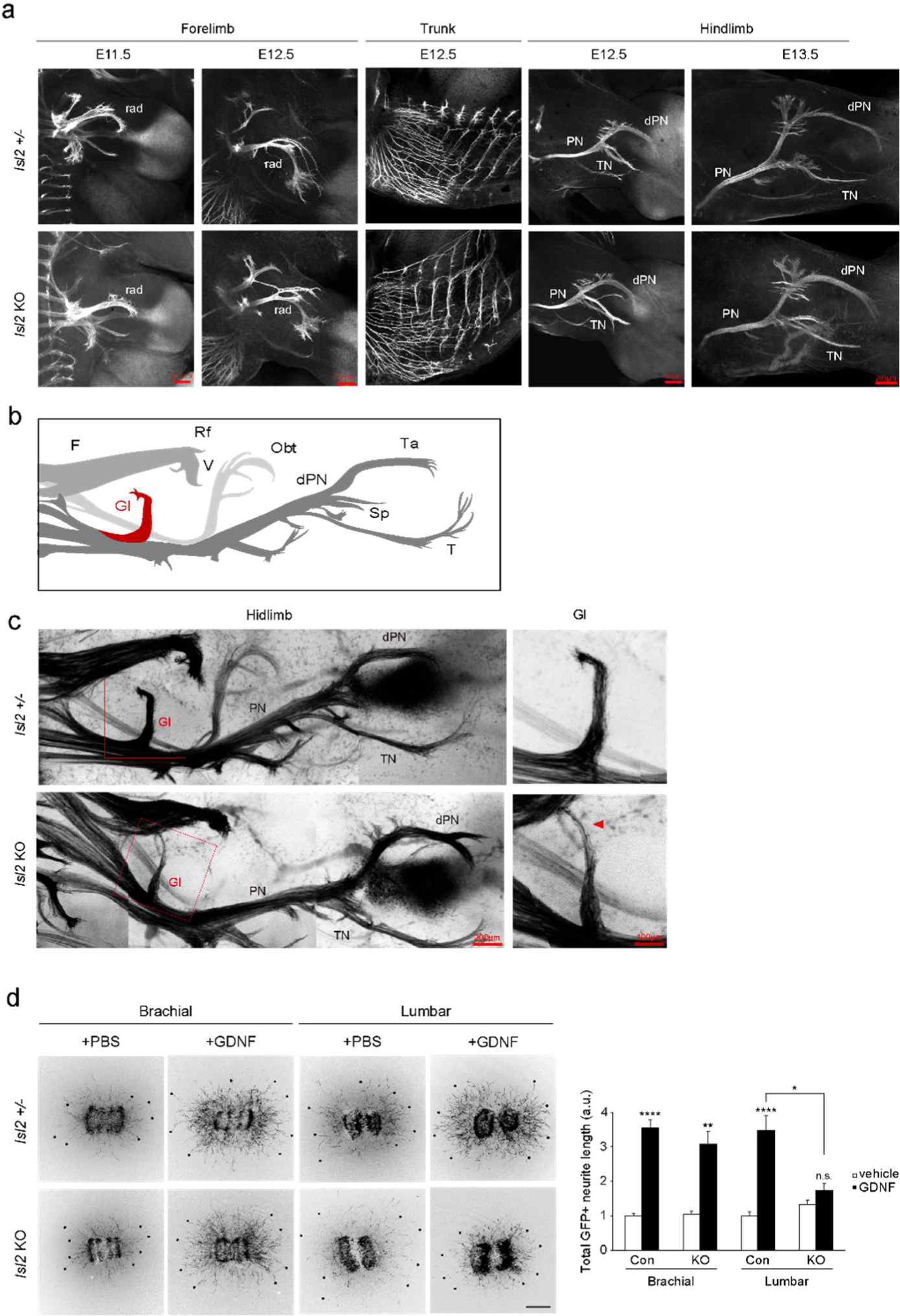
Normal innervation of the motor nerve in *Isl2* mutant hindlimbs. (a) GFP whole-mount immunostaining of forelimbs and hindlimbs from E12.5 and E13.5 *Isl2*^+/−^ and *Isl2* KO embryos. (b) Diagram of nerve trunks in the hindlimb. Hb9-driven GFP axons revealed major nerve trunks toward their target muscles. The femoral (F) nerve trunk developed branches that were targeted toward the rectus femoris (Rf) and vasti (V) muscles, and the peroneal (P) nerve trunk gave rise to branches that were targeted toward the gluteus (Gl) and tibialis anterior (Ta) muscles. (c) Hb9::GFP-labeled spinal motor nerve projection in E13.0 *Isl2*^+/−^ and *Isl2* KO embryos. dPN, deep peroneal; F, femoral; Gl, gluteus nerve; Obt, obturator; PN, peroneal nerve; rad, radial; Rf, rectus femoris; Sp, superficial peroneal; T, tibialis; Ta, tibialis anterior; V, vasti. Scale bars as indicated. (d) Representative images of motor neuronal outgrowth in explant cultures with or without GDNF in *Isl2*-deficient brachial and lumbar spinal cords. Quantification of neurite lengths (n > 3 animals per group; SEM is shown; Kruskal-Wallis test; see Supplementary Table S2 for detailed n and statistics). \**p* < 0.05, \*\**p* < 0.01, \*\*\*\**p* < 0.0001, n.s = not significant.

**Supplementary Figure 9.**
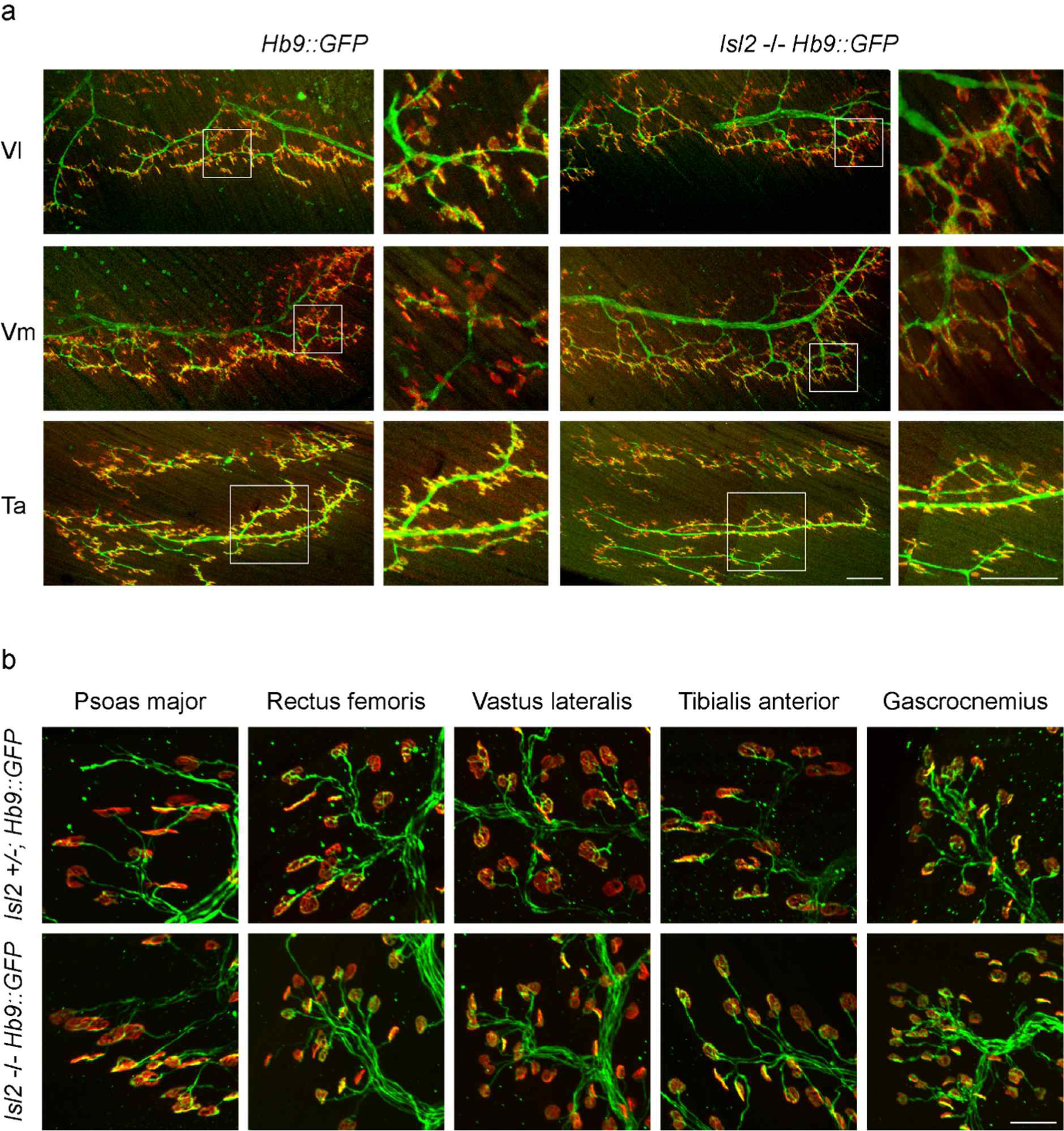
Organization of neuromuscular junctions in various hindlimb muscles of *Isl2* mutants. (a) Neuromuscular junctions in Vl, Vm, and Ta muscles of P0 control *Hb9::GFP* or *Isl2* KO; *Hb9::GFP* animals were visualized with GFP (green) and α-bungarotoxin (BTX). The boxed regions on the left are shown in higher magnification on the right. Vl, vastus lateralis; Vm, vastus medialis. Scale bars, 200 μm. (b) Neuromuscular junctions in the psoas major, rectus femoris, vastus lateralis, tibialis anterior, and gastrocnemius muscles of *Isl2^+/−^*; *Hb9::GFP* or *Isl2* KO; *Hb9::GFP* animals at P14 were visualized with GFP (green) and α-bungarotoxin (BTX). Scale bar, 50 μm.

**Supplementary Figure 10.**
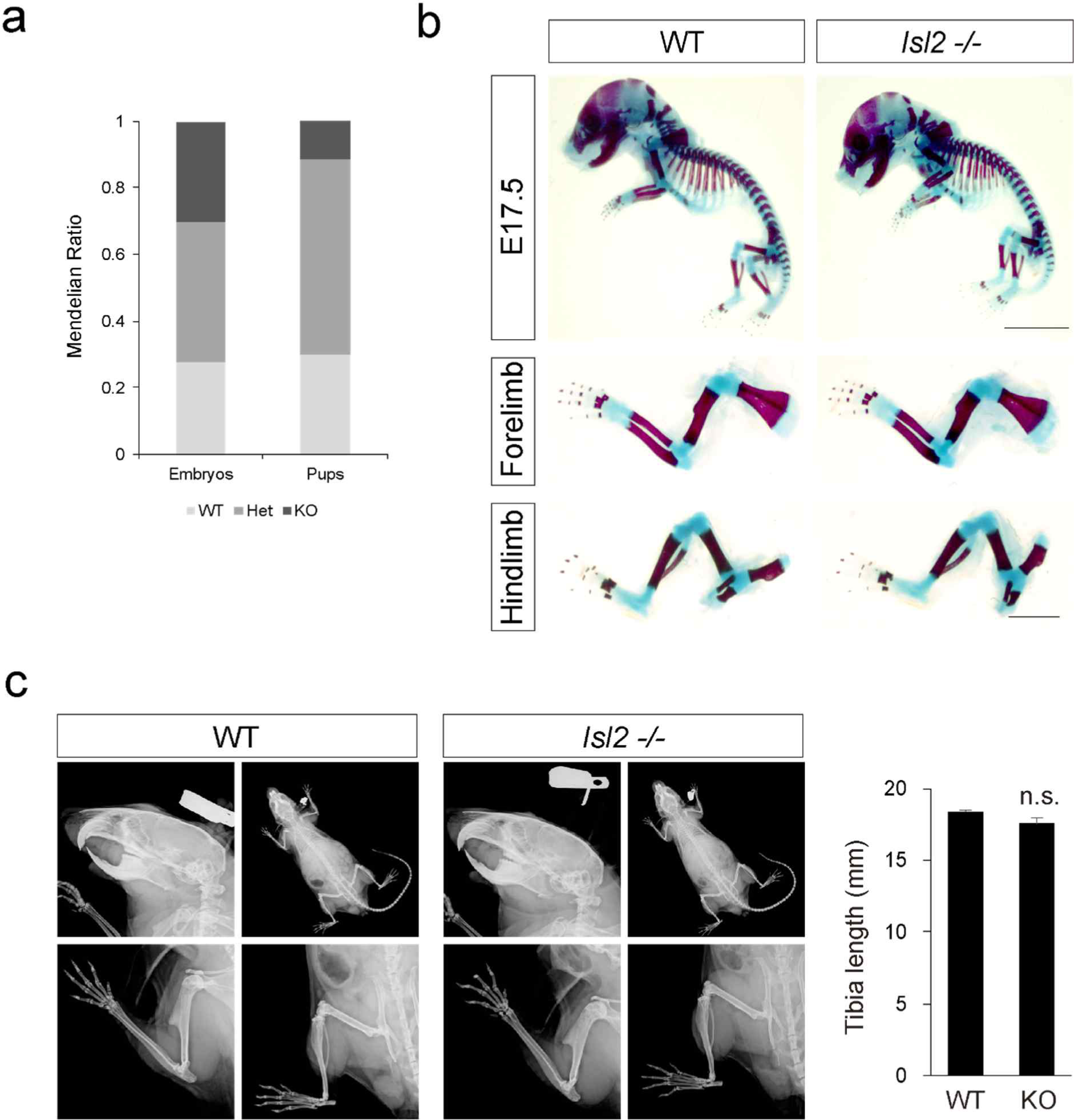
Perinatal lethality and skeletal analysis of *Isl2* mutant mice. (a) Mendelian ratios from *Isl2* heterozygous cross (n > 45 pups; n > 85, E11.5–17.5 embryos per genotype). (b) Staining with Alcian blue and Alizarin red showed no skeletal abnormality in E17.5 *Isl2*-null mice (n = 3 animals per genotype). (c) X-ray analysis showed normal bone formation in adult *Isl2*-null mice. Quantification of tibia length (left; WT, n = 5 animals; KO, n = 3 animals). SEM is shown; unpaired Student’s t-test; see Supplementary Table S2 for detailed n and statistics. n.s = not significant.

**Supplementary Table S1.**
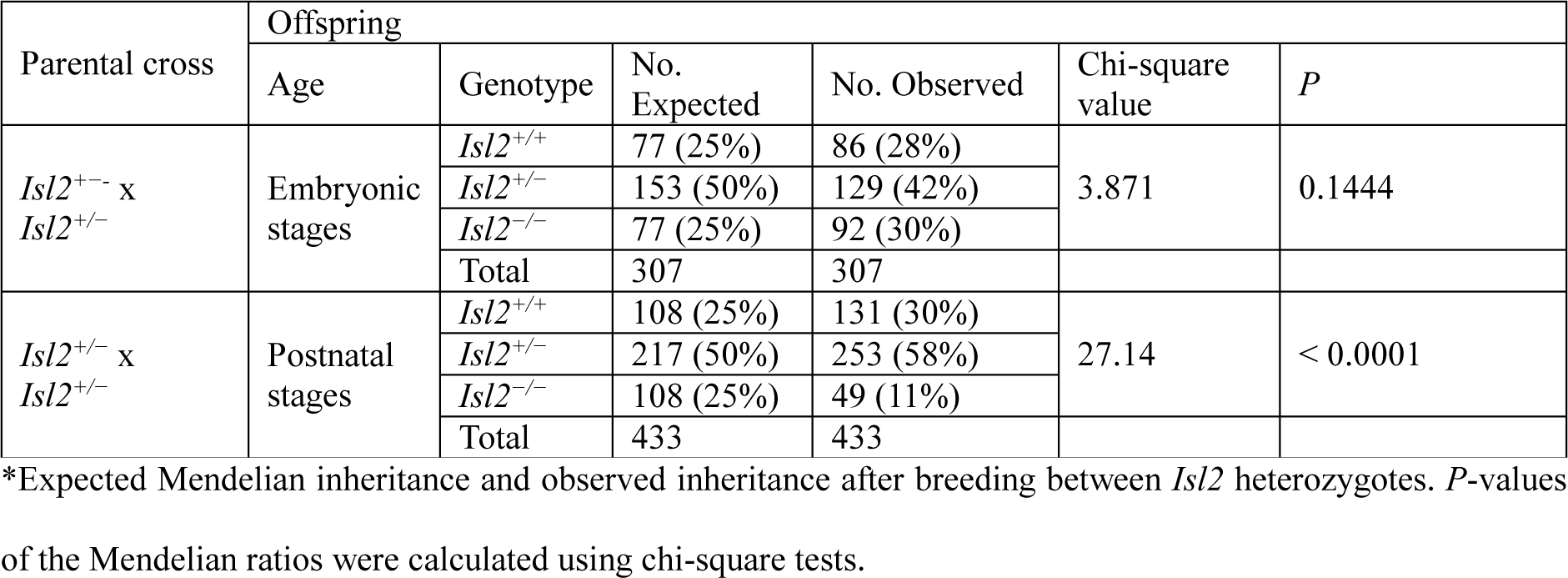
Genotype distribution of *Isl2* KO mice.

**Supplementary Table S2.**
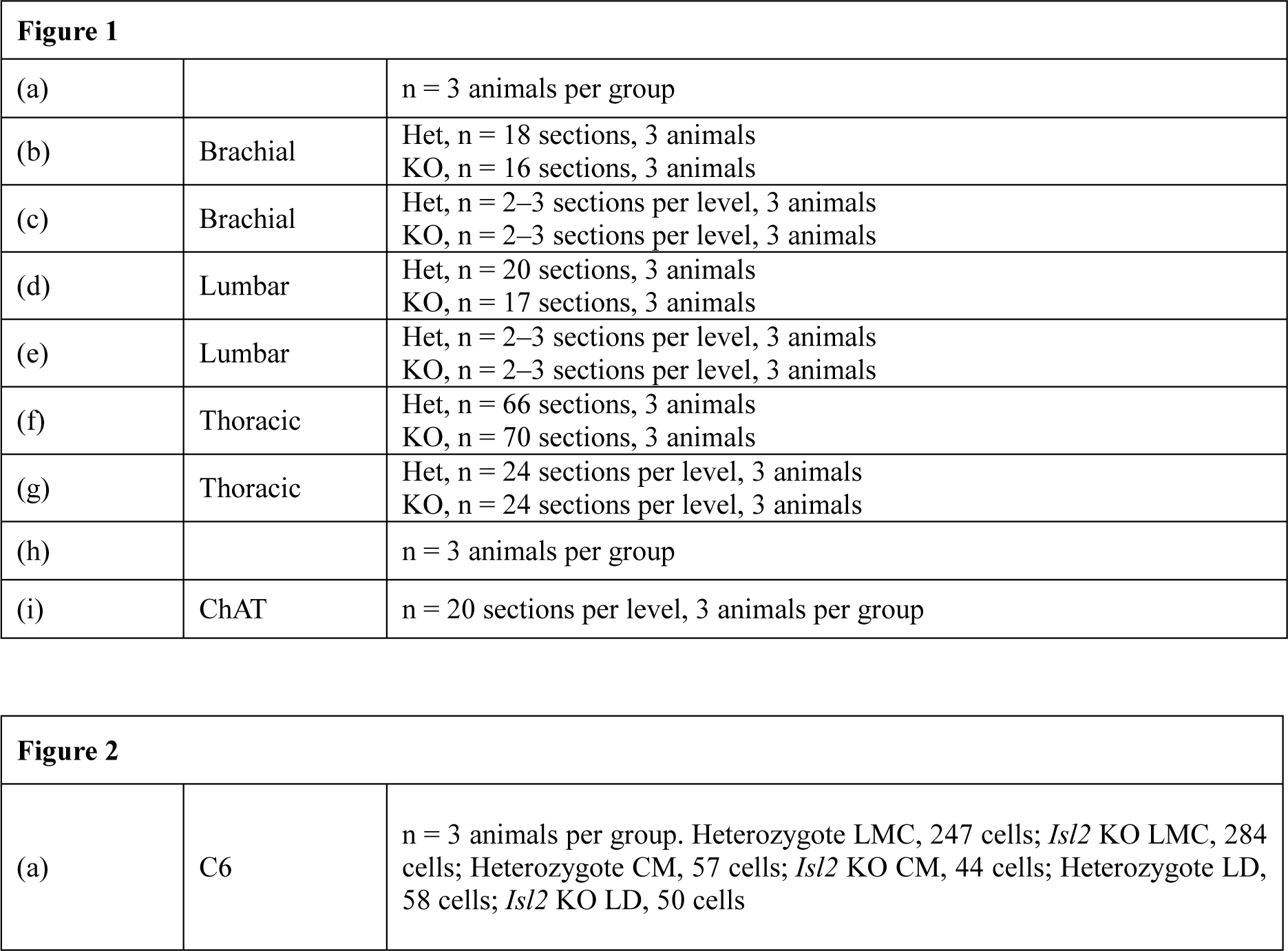
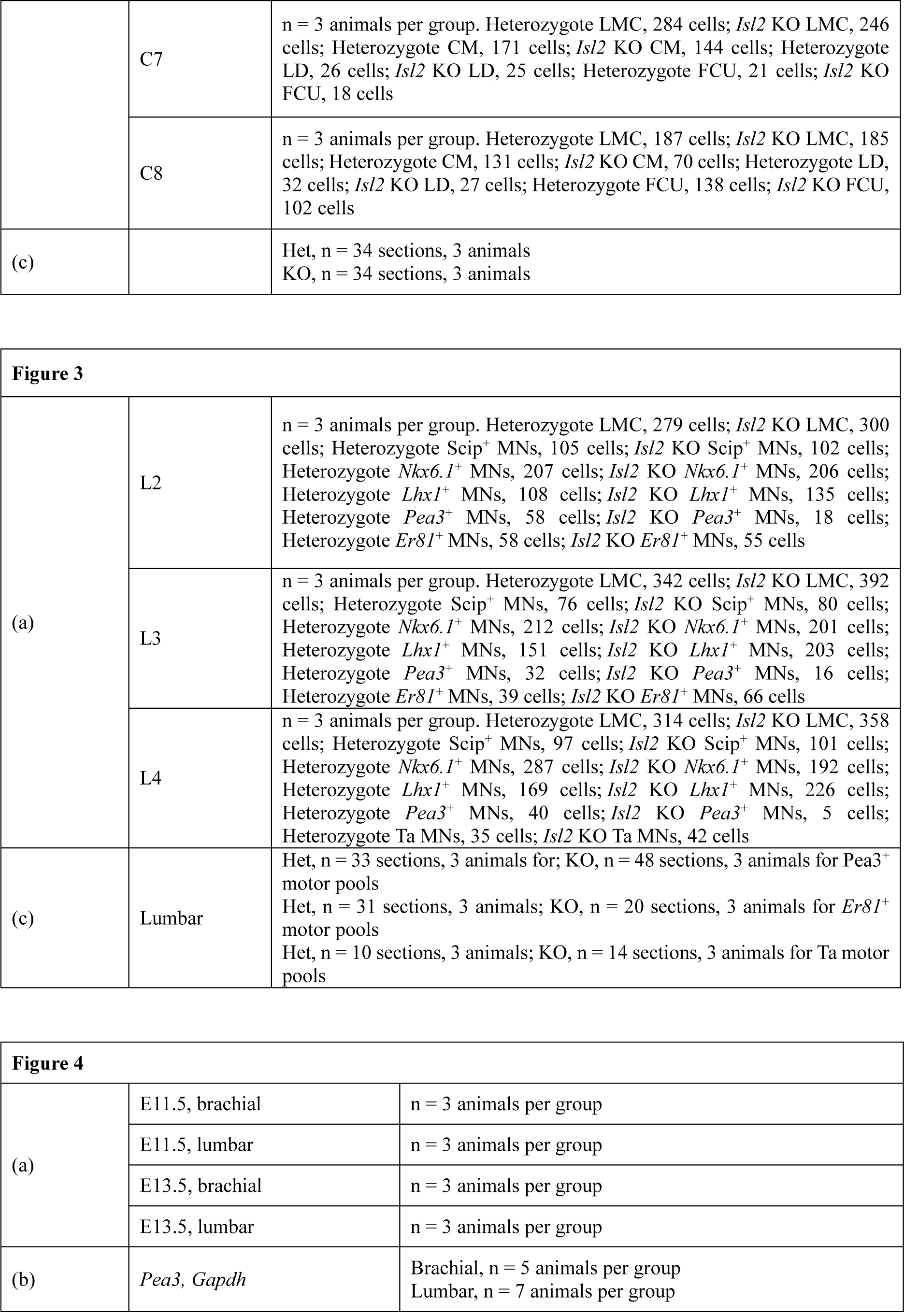
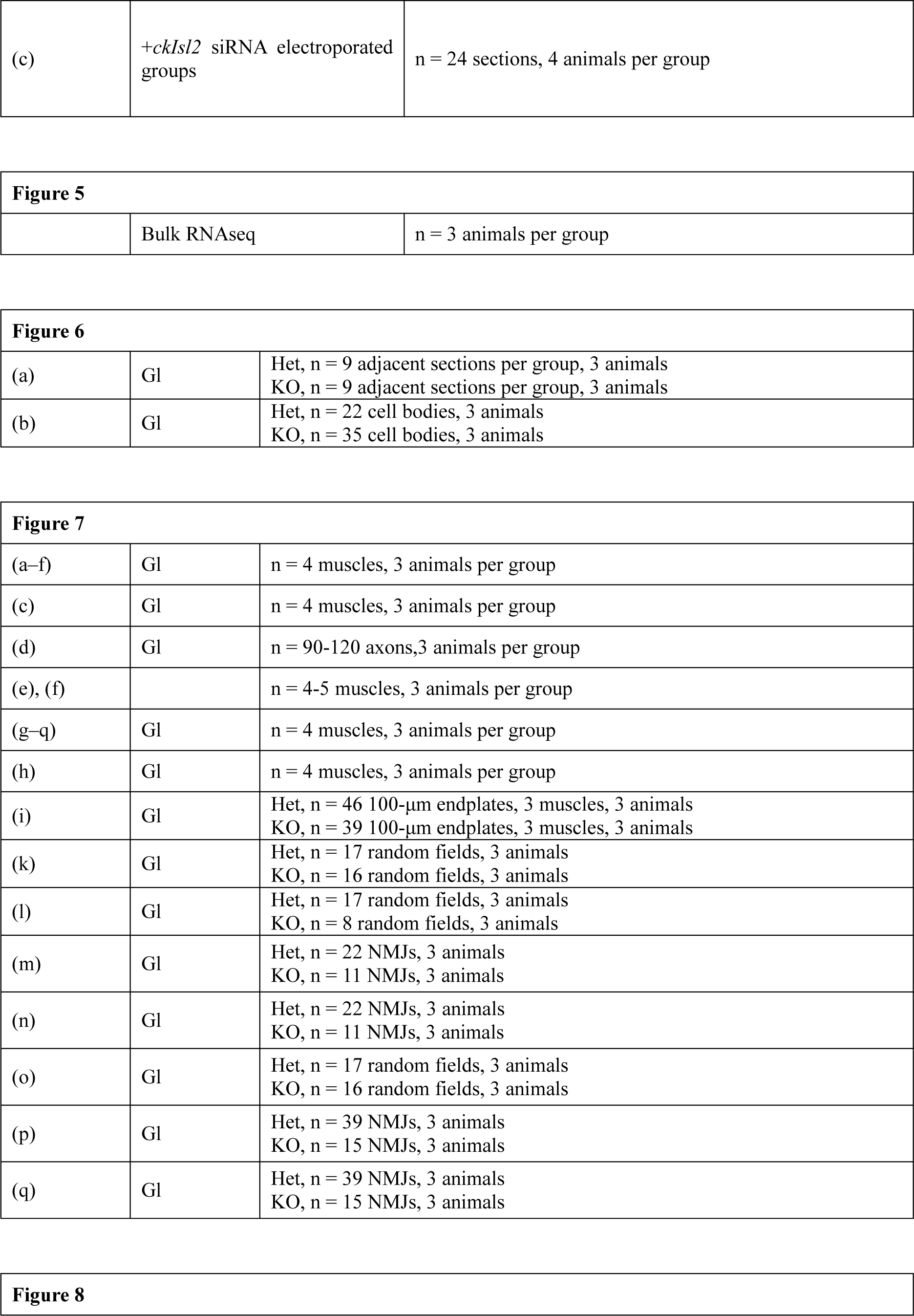
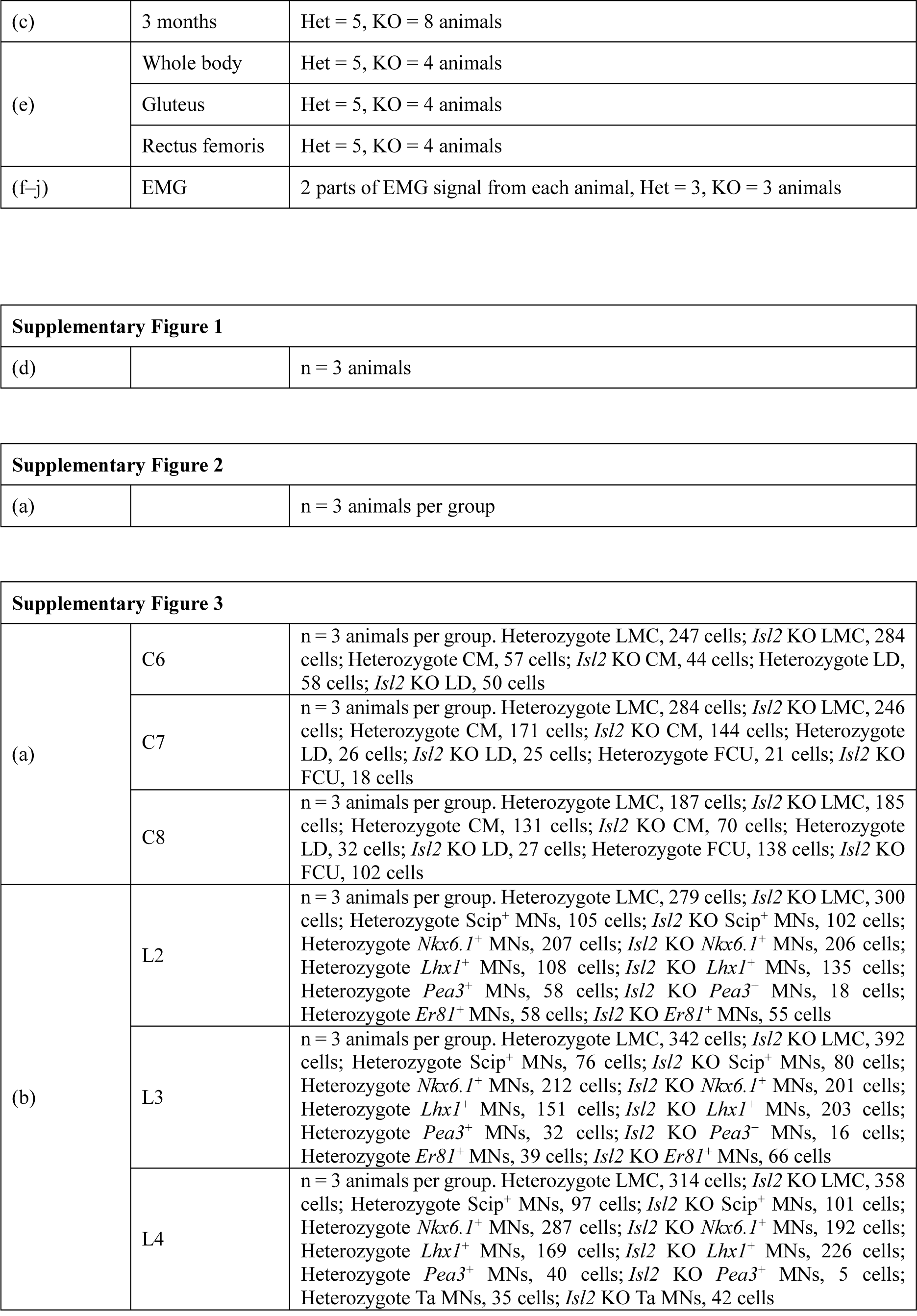
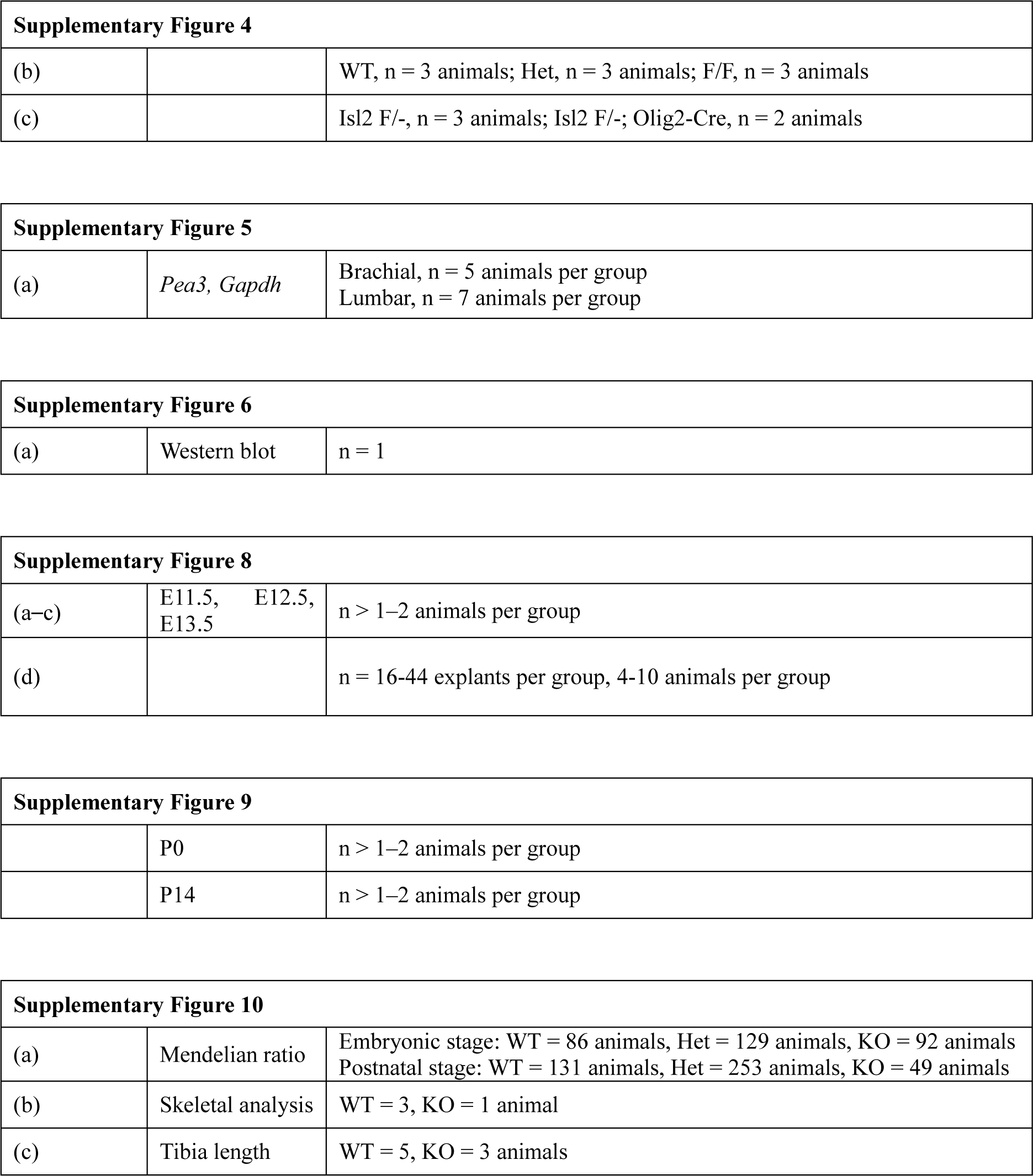
Experimental sample sizes.

**Supplementary Table S3.**
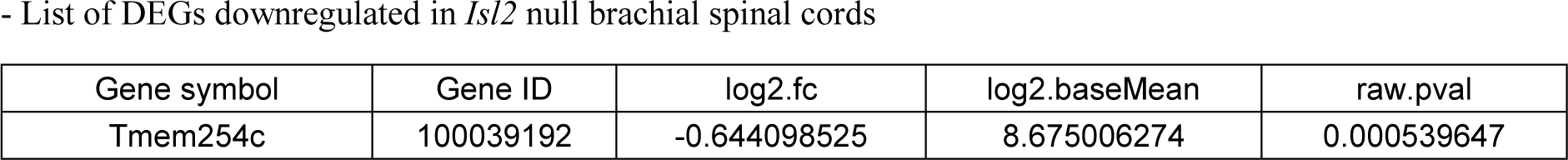
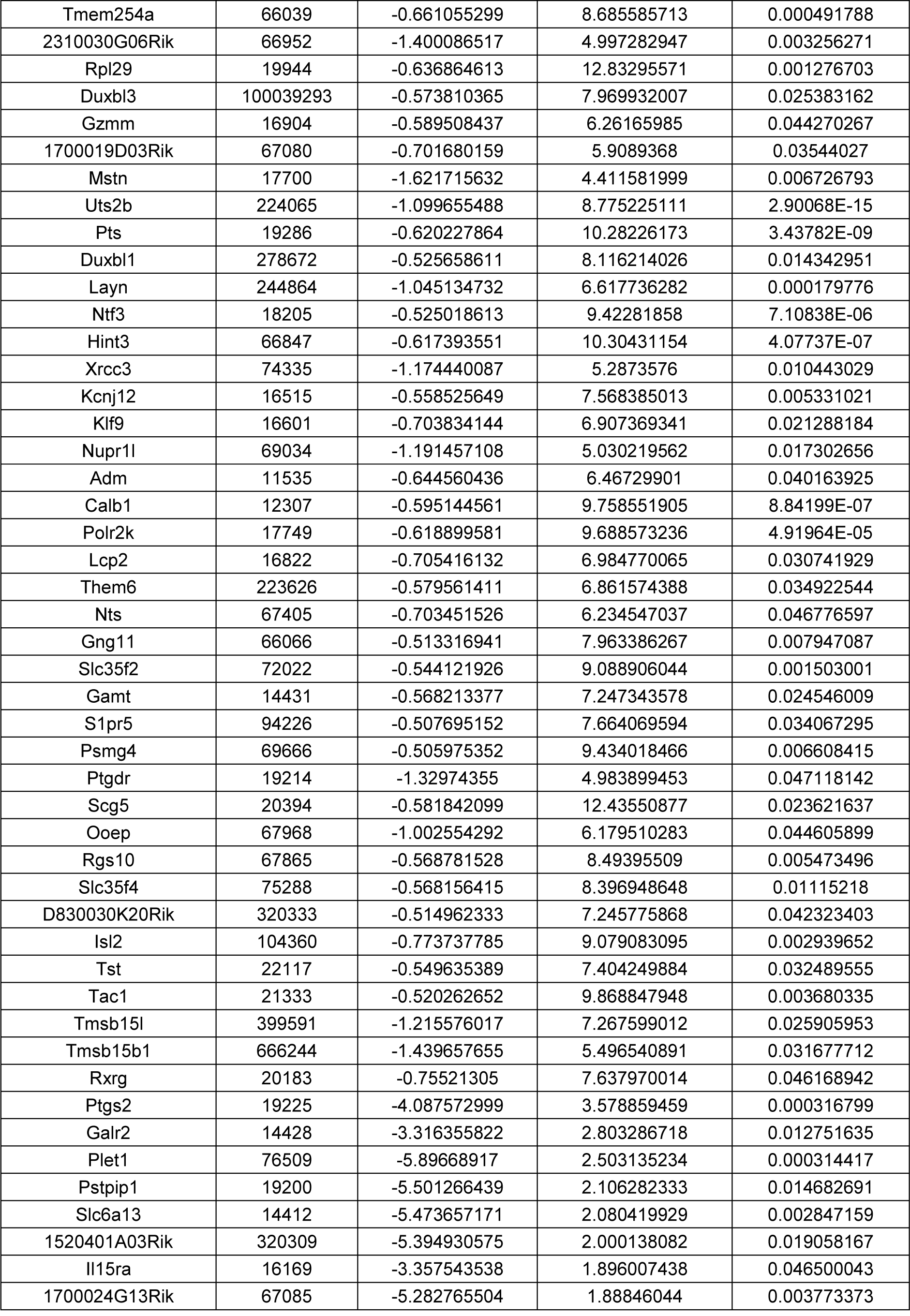
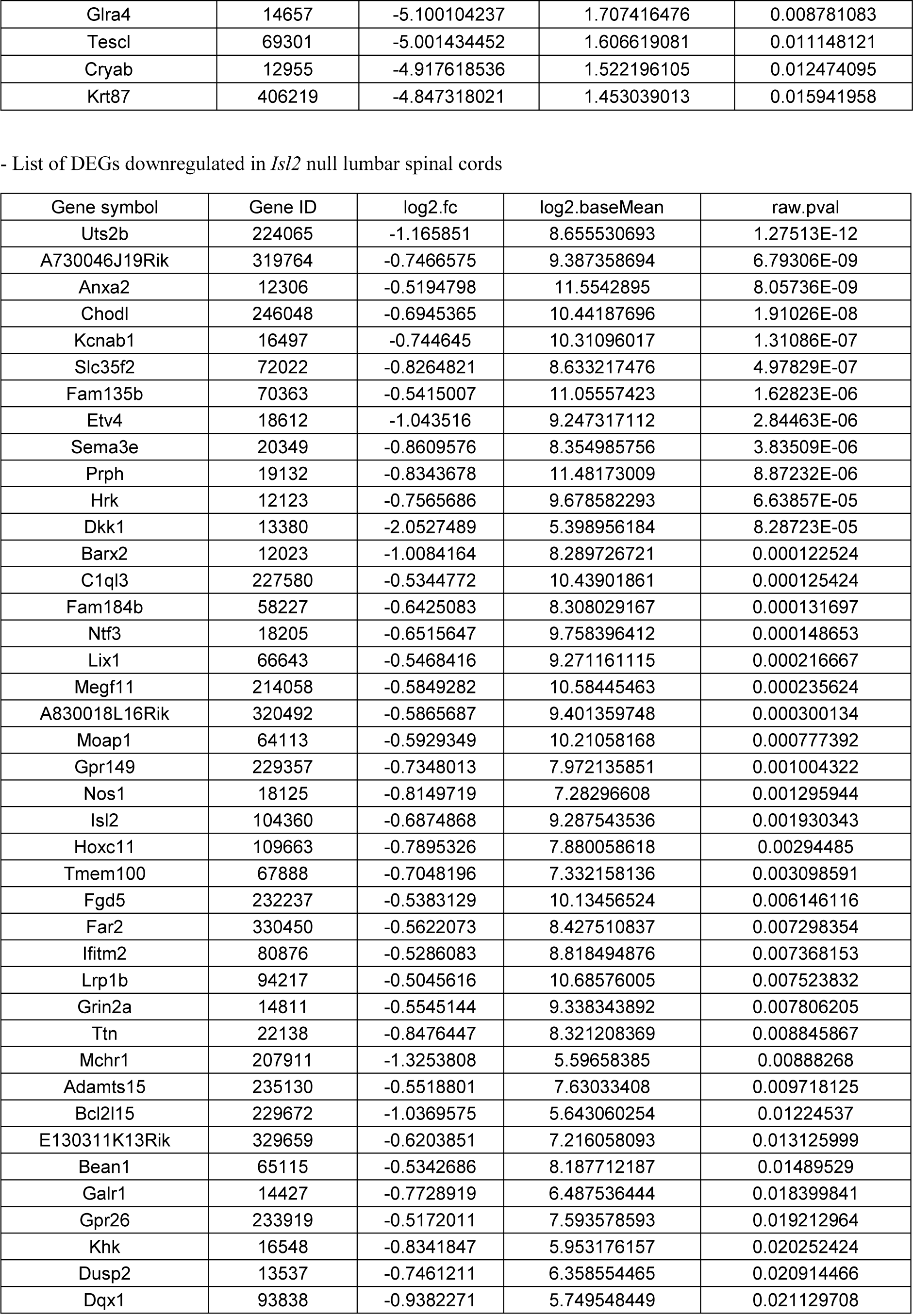
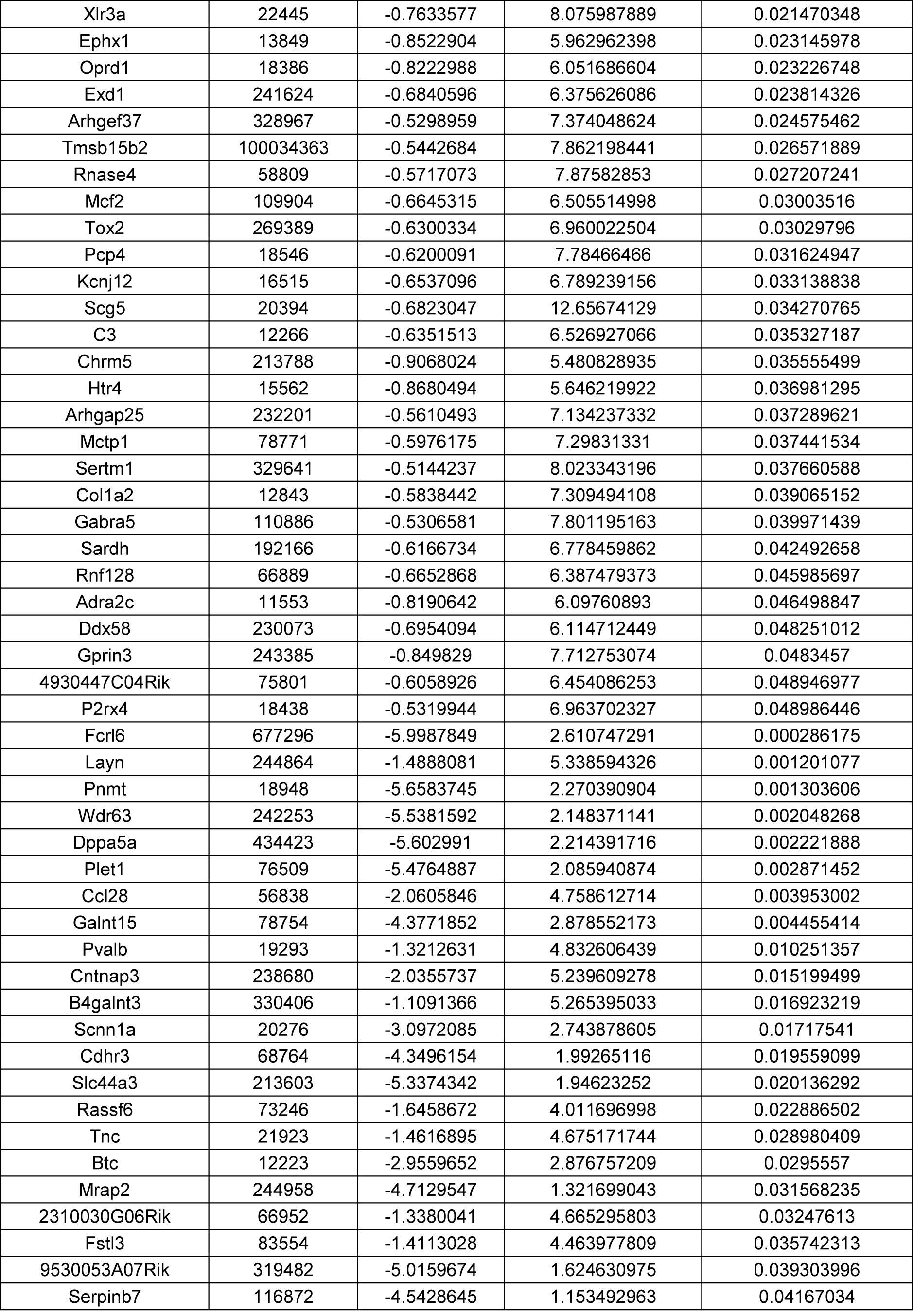
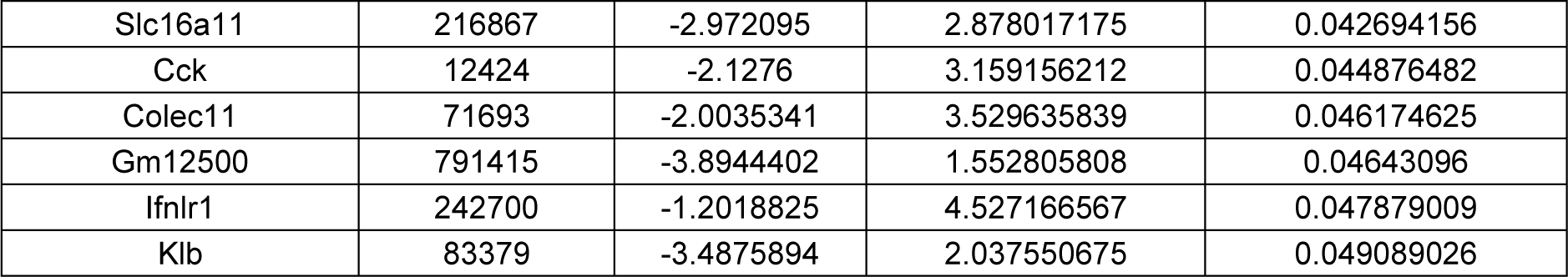
List of DEGs downregulated in *Isl2*-null brachial and lumbar spinal cords.

## Supplementary methods

### Genotyping PCR of *Isl2* flox mice

Genomic DNA was isolated from mouse ear punches digested in DirectPCR (tail) (VIAGEN) and Proteinase K (New England Biolabs). The following primers were used for genotyping PCR: 5’-TGG GAC TAC GGG GTT GTA CTT-3’ and 5’-GTT CTG GAG AGC AAG TTG GGA AT-3’ to detect a wild-type allele (274 bp) and a flox allele (410 bp).

### Immunohistochemistry and *in situ* hybridization

Immunohistochemistry was performed as described previously (Song et al., 2009). The following antibodies were used: guinea pig anti-Lhx3 (Thaler et al., 2004), rabbit anti-Foxp1 (Abcam), goat anti-Isl1 (R&D Systems), and guinea pig anti-Isl2. For *in situ* hybridization, embryonic mouse cDNA at E12.5 was used to generate riboprobes for *Isl2* using a one-step RT-PCR kit (Solgent).

### Western blot analysis

We cultured and transfected 293T cells with an expression plasmid containing N-terminal truncated chick *Isl2* (aa 38-356) with an HA tag and *Isl2* siRNA using lipofectamine (Invitrogen). After 36 hours, the cells were dissociated and lysed in lysis buffer for western blotting. Western blotting was performed as previously described(Song et al., 2009). The following antibodies were used: rat anti-α-tubulin (AbD Serotec) and mouse anti-HA (Covance).

### Skeletal staining and X-ray imaging

The skin, viscera, and adipose tissues of E17.5 mouse embryos were removed and fixed in 95% ethanol for a minimum of 5 days. Tissues were incubated in acetone for 2 days to remove fat and stained in staining solution (1 volume of 0.3% Alcian blue in 70% ethanol, 1 volume of Alizarin red in 95% ethanol, 1 volume of glacial acetic acid, and 17 volumes of 70% ethanol) for 3 days at 37°C. Stained tissues were rinsed in distilled water and cleared in 1% KOH, 20% glycerol in 1% KOH, 50% glycerol in 1% KOH, and 80% glycerol in 1% KOH, sequentially, for several days each. Cleared embryos were stored in 100% glycerol before imaging.

### Whole-mount immunostaining for axon tracing

E11.5, E12.5, and E13.5 *Hb9::GFP* embryos were dissected, and samples were immersed in 4% PFA and washed in PBS overnight. Tissues were cleared using a three-dimensional imaging kit (Binaree). Images were captured using a Zeiss confocal microscope using the ZEN software (Zeiss).

### Motor neurite outgrowth

Motor explants were dissected and isolated from E12.5 *Hb9::GFP* embryos, embedded into a collagen/Matrigel mixture (BD Sciences), and cultured in MN medium (Neurobasal medium containing B27 supplement, 2 mM L-glutamine, 25 mM L-glutamate, and 1% penicillin/streptomycin [Invitrogen]) for 30 hours (Kim et al., 2016). GDNF (200 ng/mL, R&D systems) was used to stimulate axon outgrowth. Images were captured using an LSM 700 confocal microscope (Zeiss), and neurite length was quantified using the ZEN software (Zeiss).

### Use of published datasets

For the evaluation and comparison of candidate genes downregulated in *Isl2*-deleted embryos, we compared our gene lists to that of a previously published dataset (Amin et al., 2021). The raw data of the study by Amin et al. were obtained from ArrayExpress accession: E-MTAB-10571. Reads from wild-type samples (E12 *Hb9::GFP*) were mapped to the mm10 mouse reference genome, and raw unique molecular identifiers were produced using Cell Ranger (version 3.1.0, 10x Genomics). The expression matrixes were processed in R (v.3.6.0) using Seurat (v.3.2.3). The filtering of cells and genes was performed using only cells that had a minimum of 1000 detected transcripts; cells with over 8% of mitochondrial gene read counts were excluded. Filtered gene-barcode matrixes were normalized using the *NormalizeData* function with *LogNormalize*, and the top 2000 variable genes were identified using the ‘vst’ method in *FindVariableFeatures*. Gene expression matrixes were scaled using the *ScaleData* function. We then performed principal component analysis (PCA) and UMAP using the 20 principal components. To remove the batch effect, *RunFastMNN* was run on the PCA matrix above using default parameters with SampleID as the batch key. Dimensionality reduction was performed separately for all cells and the interneuron-excluded subset. The cells were partitioned into 31 clusters, with a resolution parameter of 2. Neuron subtype identification was performed as previously described (Amin et al., 2021). Motor neuron subtypes were identified according to the expression of known marker genes. The number of cells in each neuron subtype was as follows: 19 cells in pMN, 672 cells in immature MNs, 3464 cells in postmitotic MNs, and 1135 cells in interneurons.

Further downstream analyses within LMC clusters were performed using Seurat v.4.0.2 R packages. To analyze motor pools at the L1 to L4 levels, we isolated LMC subclusters that expressed *Hoxd9*, *Hoxa10*, *Hoxc10*, *Hoxd10*, *Hoxa11*, *Hoxc11*, and *Hoxd11*. As a result, LMC clusters were further divided into seven clades composed of 276 cells: sb.LMCl.1.v, sb.LMCl.1.ta, sb.LMCl.2, sb.LMCm.1, sb.LMCm.2, and sb.LMCm.3. *Lhx1*, an LMCl marker, was enriched in sb.LMCl.1.v, sbLMCl.1.ta, and sb.LMCl.2 clades, whereas *Isl1*, an LMCm marker, was enriched in LMCm.1, LMCm.2, and LMCm.3. Visualized data were plotted using *VinPlot*, *DotPlot*, *DoHeatmap*, and *FeaturePlot* functions from the Seurat R packages.

**Video 1.** Example movie of a P14 littermate control *Isl2*^+/−^ mouse and an *Isl2* KO mouse during walking

**Video 2.** Example movie of an adult littermate *Isl2*^+/−^ and an *Isl2* KO mouse during walking

